## Supplementary material for "Single-Molecule Force Spectroscopy of Toehold-Mediated Strand Displacement": SI

### Experimental procedures and data analysis

#### Culture media

We employed LB medium (Carl Roth) and Turbo® cloning strain to culture our cells. The medium was supplemented with 100 µg/ml kanamycin.

#### Plasmid construction and cloning process

All DNA and RNA oligonucleotides were obtained from Eurofins Genomics, Ebersberg, Germany, and biomers.net GmbH. Partial toehold hairpin and trigger sequences were constructed using a combination of overlap extension PCR, restriction ligation, and blunt-end ligation. First, we amplified each part of the sequence using overlap extension PCR and annealed the resulting DNA oligos with complementary overhang sequences in Q5® Master Mix (NEB) at a calculated annealing temperature (NEB Tm calculator (https://tmcalculator.neb.com/#!/main). DNA polymerase was used to extend the 3’ ends and fill up the gaps. We then added forward and reverse primers, including restriction site sequences (EcoRI, XbaI, SpeI, and PstI), to the PCR mix and amplified the target strand. The PCR products were purified using the Monarch® PCR & DNA Cleanup Kit (NEB) and their concentration and quality were assessed using a Nanodrop 8000 spectrophotometer (Thermo Fisher).
Next, toehold hairpin and trigger sequences were ligated with vector using restriction ligation. Cloning vector plasmids were also digested with EcoRI and PstI and gel-purified to remove the digested strands. Finally, all three parts (with a ratio of inserts to vector of 1:3) were ligated using T4 ligase (NEB) following the standard protocol. The ligation products were then transformed into chemically competent cells (Turbo®, NEB) using a standard protocol. The cells were plated on LB agar plates containing 100 µg/ml kanamycin and incubated overnight at 37°C. A single colony was picked and checked using colony PCR. The selected colony was inoculated in 5 mL LB medium and incubated overnight at 37 °C. After

overnight culture, cells were collected and plasmids were purified using miniprep kits (QIAprep Spin Miniprep Kit).
Blunt-end ligation was used to optimize the toehold hairpin or trigger sequence. We amplified toehold hairpin or trigger constructs and the vector using primers that included a portion of the optimized sequence (refer to Primer List). Next, we in vitro phosphorylated the PCR products using T4 Polynucleotide Kinase (NEB) following the standard protocol. The phosphorylated PCR products were then ligated using T4 ligase at room temperature for 2 hours and digested with DpnI (NEB) using the standard protocol to remove any remaining original plasmid DNA. The final products were transformed into chemically competent cells (Turbo®, NEB). A list of all plasmids used in this study can be found in the DNA construct sequence list.

#### Cell culture

Recombinant plasmids were transformed into chemically and electrically competent *E. coli* cells: Turbo NEB (glnV44 thi-1 Δ(lac-proAB) galE15 galK16 R (zgb-210::Tn10)TetS endA1 fhuA2 Δ(mcrB-hsdSM)5, (rK–mK–) F′[traD36 proAB+ lacIq lacZΔM15]) were used for cloning using a standard protocol. All bacterial strains (Turbo NEB) were grown in LB media with appropriate antibiotics using 5 mL culture each in 5 mL centrifuge tubes at 37°C while shaking at 250 rpm. For cloning strains, a colony was picked up from an LB agar plate (Carl Roth) and inoculated into LB 5mL medium, followed by cell culture overnight using the above growth conditions.

#### DNA handle preparation and *In vitro* transcription

We PCR-amplified the DNA handle strands (545 bp) from Lambda phage DNA using modified primers (see Sequence and Primers). Specifically, the forward primer was labeled with two dT-Biotin or dT-Digoxigenin molecules at the 5' end, while the reverse primer included a stable abasic-site to preserve a single-stranded overhang for binding to the target molecule.
All *in vitro* gene transcription experiments with toehold hairpin and trigger RNAs were performed using a homemade *in vitro* transcription mix including a homemade T7 RNA polymerase. The T7 RNA polymerase with a 6xHis tag was expressed in *E. coli* BL21 DE3, followed by cell lysis using lysis buffer (1 mM Benzamidine, 1 mM PMSF, 1:2000 (1mU) dilution Turbo DNase from Ambion, 1 mg/ml Lysozyme of chicken egg white) and sonication, followed by purification using an ÄKTA pure Chromatography System. Next to the T7 RNAP, the TX mix contained transcription buffer (50 mM HEPES, 22 mM $MgCl_2$, 100 mM KCl, pH 7.8) and Murine RNase inhibitor (NEB) in a 20 µl or 100 µl reaction. Linear transcription templates for toehold hairpins and trigger RNAs were first amplified using PCR and purified using a Monarch® PCR Cleanup Kit (NEB). The concentration and quality of purified DNA templates were quantified via their 260/280 and 260/230 ratios using a Nanodrop 8000 spectrophotometer (Thermo Fisher). The molar concentration of each DNA template was calculated via:

$$\frac{Concentration\ (ng/\mu l) \times 10^6}{Molecular\ weight (g/mol)} = Concentration\ (nM) \qquad (1)$$

#### Agarose gel electrophoresis and Urea-PAGE purification

The DNA handles used for attaching the target molecule and silica beads (0.5 kb) was initially PCR amplified (primer sequence see Sequence and primers) and purified through agarose gel (2% wt, from CARL ROTH). However, we encountered a false priming issue that resulted in an additional 200 bp junk strand on the PCR product, which could significantly impact the subsequent folding process with the target molecule. The target bands were cut and purified using Gel Purification Kit from QIAGEN.
After *in vitro* transcription, the toehold hairpin and trigger RNAs were initially digested for 30 minutes using DNase I (from NEB) to remove the original DNA template. Next, we added 0.5 M EDTA to chelate the remaining $Mg^{2+}$ from the samples and denatured them at 65 °C. After denaturation, the RNA samples were purified using a 10% Urea-PAGE gel (consisting of Urea 4.8g, 40% Acryl (29:1) 2.5 ml, 30% APS 50 µL, TEMED 10 µL, and 10xTBE 1mL, all from Carl Roth). The gel electrophoresis was performed using the Owl™ gel system. Following gel electrophoresis, the target bands were cut from the gel and

the RNA was extracted using the ZR small-RNA™ PAGE Recovery Kit (from Zymo Research). The RNA concentrations were also measured using a Nanodrop 8000 spectrophotometer (from Thermo Fisher).

##### Toehold hairpin constructs preparation

After agarose and PAGE gel purification, we measured and calculated the molarity of each component, including DNA handles, toehold hairpins and adapter strand. Toehold hairpin strands and adapter strand are firstly mixed with 1:1 molarity ratio of dT-Biotin-DNA handles (40 mM) and dT- Digoxigenin-DNA handles (40 mM) respectively. Next, the samples were dried using Concentrator 5301 eppendorf and resuspended in folding buffers (1M NaCl, 50 mM HEPES, pH, 7.8 or 20 mM $MgCl_2$, 50 mM HEPES, pH 7.8). We incubated samples under different annealing temperature cycles (Tab. 1). We finally checked the folding constructs on agarose gel.

Table 1: Folding of toehold hairpin and adapter strand with DNA handles

| STEP | TEMP | TIME |
|---|---|---|
| Initial Denaturation | 70°C | 1 min |
| Folding | 68°C 30s<br>65 °C 20 min | 30 cycles |
| Final | 25°C | 5 minutes |
| Hold | 4-10°C | ∞ |

Folding of toehold hairpin-DNA handles with adapter-DNA handles

| STEP | TEMP | TIME |
|---|---|---|
| Initial Denaturation | 68°C | 1 min |
| Folding | 68°C 30s<br>63°C 20 min | 30 cycles |
| Final | 25°C | 5 minutes |
| Hold | 4-10°C | ∞ |

#### Preparation of the optical trap measurements and data analysis

##### Measurement Preparation

To minimize multi-binding of the DNA handle, we diluted the final constructs to a concentration of 0.4 nM and incubated them for 10 minutes with 1 µm-sized streptavidin-coated beads (Bangs Laboratories, Inc.) in 14 µL of running buffer (20 mM $MgCl_2$, 300 mM KCl, 50 mM HEPES, pH 7.2) at room temperature. In the meantime, we prepared the mobile phases in 500 µL of running buffer by adding an additional oxygen scavenger system (final concentrations: 26 U/ml glucose oxidase (SIGMA-ALDRICH), 17 000 U/ml catalase (SERVA), and 0.65 % glucose (SIGMA-ALDRICH)). The trigger phase included 0.1 µM of purified trigger strand, while the bead phase consisted of a mixture of streptavidin-coated beads and anti-digoxigenin-coated beads incubated together in 300 µL of the same scavenger system. All the phases were added to the syringe pump of the C-Trap® Optical Tweezers – Fluorescence & Label-free Microscopy. We used a commercial microfluidic chip with multiple inlets that allowed us to generate a laminar flow to separate different phases (see Fig. 1B). In the bead channel, we trapped the two different kinds of beads, one in the fixed beam and the other one in the mobile beam. In the buffer channel, the molecular construct was tethered between the two beads by mobilizing one trap towards the fixed trap, resulting in the close proximity of the bead surfaces and the formation of a dumbbell-shaped conformation, known as the ‘dumbbell assay’ (See Fig. 1A). We maintained the laminar flow of buffer and trigger phase by applying ~ 0.35 bar to the syringes, leading to a flow velocity of ~ 20 µm/s during the experiment to inhibit trigger diffusion into the buffer channel even for measurement times of up to an hour. In a typical experiment, designed to measure the invasion time, as shown in Figure 2C and D, we initially established the tether between two beads and then transitioned to the trigger channel for the specific purpose of measuring trigger binding and strand invasion events exclusively. In all passive mode experiments, after binding the trigger strand in the trigger channel, we promptly relocated the beads back to the buffer channel to conduct pulling cycles and passive mode measurements. This step was taken to mitigate the potential influence of other trigger strands competing with the bound trigger strand and unspecific binding of trigger strands to unfolded parts of the toehold hairpin. The trap stiffness used in all measurements was between 0.25 pN/nm and 0.40 pN/nm. The sampling rate was 78.125 kHz and was down sampled by a factor of 3 for data analysis except for the invasion time determination and autocorrelation analysis used for data shown in Fig. 2C and D. Measurement temperatures were ~ 25°C.

**Stretch and relax cycles**

Without trigger strand:

Once the tether was formed, we performed several stretch-and-relax cycles at a constant velocity of 0.2 µm/s by moving the traps apart to a distance where the toehold hairpin was fully unfolded and then reducing the trap distance to allow the hairpin to refold. Force (F) versus extension (e) traces were used to determine the unfolding intermediates of the toehold hairpin including the associated contour length gains for calculating the opened base pairs.

Modeling polymer elasticity in stretch and relax cycles:

The force-extension curves for stretching of the dsDNA linkers only (toehold hairpin in the folded state) can be modeled by using the extensible worm-like-chain (eWLC) model[1,2]

$$F_{eWLC}(e) = \frac{k_B T}{p_{dsDNA}}\left[\frac{1}{4}\left(1 - \frac{e}{L_{dsDNA,linker}} + \frac{F}{K}\right)^{-2} - \frac{1}{4} + \frac{e}{L_{dsDNA,linker}} - \frac{F}{K}\right] \quad (2)$$

where $k_{\mathrm{B}}T$ is the thermal energy, $p_{\mathrm{ds}DNA}$ the dsDNA linker persistence length, $L_{\mathrm{ds}DNA,\mathrm{linker}}$ the dsDNA linker contour length and $K$ the elastic stretch modulus. If parts of the DNA/RNA toehold hairpin are unfolded, the elastic behavior can be described by the eWLC in series with a standard WLC[1]

$$F_{WLC}(e) = \frac{k_B T}{p_{ssDNA/ssRNA}}\left[\frac{1}{4}\left(1 - \frac{e}{L_{ssDNA/ssRNA}}\right)^{-2} - \frac{1}{4} + \frac{e}{L_{ssDNA/ssRNA}}\right] \quad (3)$$

where $p_{\mathrm{ssDNA/ssRNA}}$ is the persistence length of the unfolded single-stranded DNA/RNA toehold hairpin and $L_{\mathrm{ssDNA/ssRNA}}$ the contour length of the unfolded single-stranded DNA/RNA toehold hairpin. The standard WLC only takes the entropic effects into account. The fits that include unfolded ssDNA/ssRNA ($F_{eWLC}$ in series with $F_{WLC}$) had a fixed $L_{\mathrm{ds}DNA,\mathrm{linker}}$, $p_{\mathrm{ds}DNA}$ and $K$ obtained from the previous fit of the folded state. Also, a fixed $p_{\mathrm{ssDNA}}$ of 1.0 nm and $p_{\mathrm{ssRNA}}$ of 0.9 nm was used. The number of unfolded nucleotides based on the increase in contour length was calculated as follows: First, the conversion ratio of unfolded contour length and unfolded nucleotides was calculated considering the effect of the diameter of the nucleic acid and assuming the first base pair to be already opened because of fraying[3]. Second, the unfolded contour length of each intermediate state was divided by this ratio and this value was added to the two frayed nucleotides to yield the number of unfolded nucleotides for each intermediate.

With trigger strand:

In an invasion process as shown in Fig. S3B from the toehold-bound (TB) to the fully invaded state (FI) the extension increases. This can be explained by the partial opening of the toehold hairpin. The additional extension gain is comprised to one part by the double-stranded invader-target complex (invader- 5' toehold hairpin) and to the other part by the single-stranded incumbent (3' toehold hairpin) (see schematics in Fig. S3). Since there is an increase of single-stranded as well as double-stranded extension, the procedure of fitting the force-extension trace with a series of the eWLC and WLC keeping the double-stranded part $L_{\mathrm{ds}DNA}$ fixed is no longer feasible. Since the increase in the number of additional single-stranded nucleotides and double-stranded base pairs are the same in an invasion process, one can express the additional double-stranded contour length $L_{\mathrm{ds,bm}}$ by the additional single-stranded contour length $L_{\mathrm{ss,bm}}$. Therefore, one can avoid an additional free fitting parameter. Note that a fixed constant ratio of $L_{\mathrm{ss,bm/nt}}$ and $L_{\mathrm{ds,bm/bp}}$ is needed to link the two variables. Here, we used a single-stranded contour length per nucleotide of 0.59 nm/nt for DNA and 0.6 nm/nt for RNA. For the double-stranded contour length per base pair, we used 0.34 nm/bp (DNA-DNA)[4,5], 0.30 nm/bp (DNA-RNA)[4] and 0.28 nm/bp (RNA-RNA)[6]. These values are used for calculating the theoretical values of opened contour length. The total unfolded contour length was in good agreement with an unfolding of 110 nucleotides assuming a conversion of 0.6 nm/nt for ssRNA and 0.59 for nm/nt for ssDNA and taking the diameter of the nucleic acid into account. To fit for example the FI state of RRp2, we use again the series of the eWLC and WLC but adjusted for the invasion process where we adapt it to

$$\begin{gathered} L_{ssDNA/ssRNA} \rightarrow L_{ss,bm} \\ L_{dsDNA,linker} \rightarrow L_{dsDNA,linker} + \frac{L_{ds,bm/bp}}{L_{ss,bm/nt}} \cdot L_{ss,bm} \end{gathered} \tag{4}$$

and where $L_{\mathrm{dsDNA,linker}}$ , $p_{\mathrm{dsDNA,linker}}$ and $K$ are fixed and have the values obtained from a fit of the TB state using the eWLC. The only free fitting parameter is $L_{\mathrm{ss,bm}}$.

**Passive-mode traces**

Without trigger strand:

We also carried out so-called passive mode experiments where we kept the distance between the lasers constant while observing the fluctuations of the molecule through its intermediate states (Fig. S2). Assignment and coloring of the states was done using hidden-Markov-modeling (HMM)[7]. In the case of passive mode experiments, the already state-assigned force data was transformed to a contour length by inverting eq. S2 and S3, using the elastic parameters obtained in the stretch-and-relax cycles[8] and the same two steps as in the stretch and relax cycles were performed to calculate the number of unfolded nucleotides for each intermediate. This approach further allowed us to extract free energies and kinetic rates of these states, as demonstrated for example in Figure 2C, D, Figure 3C, D, and Figure 5C, D.

With trigger strand:

In the case of an invasion/re-invasion transition as in Fig. 3C, the state assignment is done in the same way as for unfolding/refolding transitions. To obtain the opened contour length we again used the contour length transformation in previous work[8] but now with the eWLC and WLC that have the adjusted parameters from Eq. 4.

**Extracting free energies**

Without trigger strand:

For calculating the free energies at zero load for each constant distance step, we used a model introduced previously for protein folding under force that takes all energetic contributions from the dumbbell assay (as shown in Fig. 1 and SI 'dumbbell assay') into account[9]. The system bead-dsDNA-ssDNA/ssRNA-dsDNA-bead is simplified to an equivalent bead-dsDNA-ssDNA/ssRNA system with an effective trap stiffness $k_{eff}^{-1} = k_1^{-1} + k_2^{-1}$, a combined contour length of both dsDNA linkers and an effective bead deflection $x_{eff} = |x_1| + |x_2|$. Since the passive mode traces are not constant force measurements, the change in extension and force in folding/refolding transitions has to be considered in the free energy calculations. The energy $G_i(F_i)$ that is stored in the bead-dsDNA-ssDNA/ssRNA system at force $F_i$ can be divided into the energy stored in the displacement of the beads $G_i^{beads}(F_i)$ and the energy stored in the stretched dsDNA linkers $G_i^{dsDNA,linker}(F_i)$ and the unfolded ssDNA/ssRNA $G_i^{ssDNA/ssRNA}(F_i)$ as well as the free energy $\Delta G_{0,i}^{DNA/RNA}(F_i)$ of the DNA/RNA toehold hairpin in state *i*: $G_i(F_i) = \Delta G_{0,i}^{DNA/RNA}(F_i) + G_i^{beads}(F_i) + G_i^{dsDNA,linker}(F_i) + G_i^{ssDNA/ssRNA}(F_i)$ . The individual terms can be calculated as

$$G^{beads}(F) = \frac{1}{2} k_{eff}^{-1} F^2 \quad (5)$$

$$G^{dsDNA,linker}(F) = \int_0^{e_{eWLC}(F)} F_{eWLC}(e')de' \quad (6)$$

$$G^{ssDNA/ssRNA}(F) = \int_0^{e_{WLC}(F)} F_{WLC}(e')de'. \quad (7)$$

The energy difference between two states *i,j* at forces $F_i$ and $F_j$ is then given by

$$\Delta G_{ij}(F_i, F_j) = G_j(F_j) - G_i(F_i) = -k_B T \cdot ln\left(\frac{P_j(F_j)}{P_i(F_i)}\right) \quad (8)$$

using the Boltzmann equation. This allows to calculate free energy differences between different DNA/RNA toehold hairpin intermediate states at zero load as

$$\Delta G_{0,ij} = -k_B T \cdot ln\left(\frac{P_j(F_j)}{P_i(F_i)}\right) - \Delta G^{beads}(F_i, F_j) - \Delta G^{dsDNA,linker}(F_i, F_j) - \Delta G^{ssDNA/ssRNA}(F_i, F_j). \quad (9)$$

The HMM assigned state probabilities were used to obtain $P_i(F_i)$, and $P_j(F_j)$.

With trigger strand:

For invasion/re-invasion processes, the increase in extension is again divided into single-stranded and double-stranded parts. Therefore, one has to adapt the parameters

$$\Delta G^{ssDNA/ssRNA} \to \Delta G^{ss,bm}$$
$$\Delta G^{dsDNA,linker} \to \Delta G^{dsDNA,linker+ds,bm} \quad (10)$$

where the contour length in the WLC integrals is adjusted as in S4.

**Model for transition rate-extrapolation**

Without trigger strand:
From the HMM assigned states in the passive mode traces, the dwell time distributions were plotted as normalized integrated histograms (see Fig. S2E and F). They were fitted by a single exponential function that accounts for the experimental time frame and the finite time resolution[10] yielding the rate constants $k_{ij}(F_i)$ at the force $F_i$ of the starting state $i$ of the transition. Since a folding/refolding transition is characterized by step-by-step zipping/unzipping events upon force exertion, the contour length increase is a well-defined reaction coordinate. The transition state position depends on the applied force and a simple Bell model is not applicable here. We adapted a model for folding/refolding of globular proteins[11] and coiled coils[9] under force for folding/re-folding of nucleic acids that takes energy changes of dsDNA linkers, springs and unfolded ssDNA/ssRNA in an energy barrier $\Delta G_{iT}^{\#}$ into account. The force dependent rates are then described by

$$k_{ij}(F) = k_{0,i}\exp\left(-\Delta G_{iT}^{\#}(F_i = F, F_T)/k_B T\right) \quad (11)$$

with $k_{0,i}$ the folding rate constant at zero load as a fit parameter (y intercept in Fig 3D). The force dependent energy difference $\Delta G_{iT}^{\#}$ from state *i* to the transition state *T* is defined as

$$\Delta G_{iT}^{\#}(F_i, F_T) = \Delta G_{iT}^{beads}(F_i, F_T) + \Delta G_{iT}^{dsDNA,linkers}(F_i, F_T) + \Delta G_{iT}^{ssDNA/ssRNA}(F_i, F_T) \quad (12)$$

with $F_T$ the force acting on the transition state *T* between two intermediate toehold hairpin states *i* and *j*. The second fit parameter defining the slope of the fits in Fig. 3D is the contour length difference $\Delta L_{iT}^{\#}$ from state *i* to the transition state *T*.

With trigger strand:
Analogously to the previous section of extracting free energies with trigger strand, the free energy contributions of the double-stranded and single-stranded parts in the invasion/re-invasion transition have to be adapted as in S10.

**Invasion time determination and autocorrelation analysis**

To obtain the invasion time $\tau$ of the invasion transitions for DNA Fig. 2C and D, the transients were fitted with an exponential function:

$$F(t) = y_0 + A\ exp(-(t_0 - t)/\tau) \quad (13)$$

For RNA (Fig. 2C and D), transient durations were significantly longer and transition times could be readily obtained by visual inspection of the traces.

To determine the system response time (gray symbols and fit in Fig. 2D), we computed the autocorrelation of the noise in regions before the transient had occurred as described in previous work[12].

**Strand displacement and mean first passage time for a 1D random walk**

Master equation:
The strand displacement process can be treated by a master equation for a 1D random walk among N+1 states (from state 0 to state N). Each state j can transition to its neighboring states j-1 and j+1 with respective rates $k_j^-$ and $k_j^+$.
The time evolution of the probability $P(j,t)$ of being in state *j* at time *t* is given by:

$$\frac{dP(j,t)}{dt} = k_{j-1}^{+}P(j-1,t) + k_{j+1}^{-}P(j+1,t) - \left(k_j^{-} + k_j^{+}\right)P(j,t) \quad (14)$$

The mean first passage time (MFPT) to reach the absorbing boundary at N - when state 0 is the starting state - is given by the following expression (solution of the backward Kolmogorov equation):[13,14]

$$T_0 = \sum_{m=0}^{N-1}\sum_{l=0}^{m}\frac{1}{k_l^{+}}\prod_{j=l+1}^{m}\frac{k_j^{-}}{k_j^{+}}$$

(15)
The transition rates between states *j* and *j*+1 are related to the energies $\epsilon_j$, $\epsilon_{j+1}$ via:

$$\frac{k_j^{+}}{k_{j+1}^{-}} = e^{(-\beta(\epsilon_{j+1}-\epsilon_j))} = e^{(-\beta\Delta\epsilon_j)} =: \omega_j \quad (16)$$

where $\beta = (k_B T)^{-1}$. We now assume that all energy differences are the same and thus also all $\omega_j$ have the same value $\omega$.

We make the additional assumption that all individual forward rates are the same, i.e., $r := k_j^+$ for all j. Then $r = k_j^+ = \omega k_j^-$. The expression for $T_0$ then turns into

$$T_0 = \sum_{m=0}^{N-1} \sum_{l=0}^{m} \frac{1}{r} \prod_{j=l+1}^{m} \frac{1}{\omega}$$

(17)

When all energies are same:
When all energies are equal, we get $\omega = 1$ and therefore:

$$T_0 = \sum_{m=0}^{N-1} \sum_{l=0}^{m} \frac{1}{r} \prod_{j=l+1}^{m} 1 = \frac{1}{r} \sum_{m=0}^{N-1} \sum_{l=0}^{m} 1 = \frac{1}{r} \sum_{m=0}^{N-1} (m+1) = \frac{1}{r} \frac{N(N+1)}{2}$$

(18)
i.e., $T_0 \approx N^2/2 \times \tau_0$ with the 'step time' $\tau_0 = \frac{1}{r}$. In this case, we obtain the expected diffusive scaling ~ $N^2$ of the process.

Energies differences are the same, but non-zero ($\omega > 1$):
When the strand displacement is biased due to an energy difference $\Delta\epsilon_j < 0$, we get $\omega > 1$ and the forward rates are correspondingly sped up compared to the backward rates. In this case we rewrite $T_0$ in the following way:

$$T_0 = \frac{1}{r} \sum_{m=0}^{N-1} \sum_{l=0}^{m} \frac{1}{\omega^{m-l}} = \frac{1}{r} \sum_{m=0}^{N-1} \frac{\omega - (1/\omega)^m}{\omega - 1}$$

(19)
which results in

$$T_0 = \frac{1}{r} \frac{\omega}{(\omega-1)^2} [N\omega - N - 1 + (1/\omega)^N] \quad (20)$$

When N is large:
When $\omega > 1$ and $N$ is large enough, we can neglect the term $(\frac{1}{\omega})^N$. The expression for the MFPT then becomes

$$T_0 \approx \frac{1}{r} \frac{\omega}{(\omega-1)^2} [N\omega - N - 1] \approx \frac{1}{r} \frac{\omega}{(\omega-1)^2} N(\omega - 1) = \frac{N}{r} \frac{\omega}{\omega - 1}$$

(21)
We see that already under this moderate assumption $T_0 \sim N$. If we naively just determine $T_0$ and divide it by $N$, the result will be

$$T_0/N \approx \frac{1}{r} \frac{\omega}{\omega-1} \quad (22)$$

When $\omega$ is large:
Now when $\omega \gg 1$, we will get from Eq. 20

$$T_0 \approx \frac{1}{r} \frac{\omega}{\omega^2} \times N\omega = \frac{N}{r} \quad (23)$$

or directly from the Eq. 22 above. In this case we simply have $T_0/N=1/r$, which we can identify with step time $\tau_0$.

What is $\omega$ under force:
In a very naive picture, the energy of state $j$ at position $x_j = x_0 + j \cdot \Delta x$ will be lowered by a force $f$ like:

$$\epsilon_{j+1}(f) = \epsilon_{j+1}(0) - f \cdot (j+1)\Delta x = \epsilon_j(f) - f \cdot \Delta x \quad (24)$$

When all energies in the absence of force are equal, then $\Delta\epsilon(f) = -f \cdot \Delta x$ and therefore $\omega = e^{(\beta f \cdot \Delta x)}$. When $\Delta x$ is ≈ 0.7 nm, then at $f$ = 10 pN the term $f \cdot \Delta x \approx 1.7\, k_B T$ at experimental conditions and thus $\omega \approx 5.5$. In this case we would have scaling of $T_0 \sim N$ and $T_0/N$ would be ≈ 22 % larger than $\tau_0$.

### Simulation protocols

We created the DNA handle and toehold hairpin structure in OxDNA using oxview[15] and we use oxDNA2 and oxRNA2 version of the model for DNA simulations[16,17], with sequence dependent parametrization. Molecular dynamics (MD) simulations are utilized for relaxation of DNA constructs and pulling simulations.

##### Kinetic simulations

Molecular dynamics simulations of the DD and RR system were performed using the oxDNA2 / oxRNA2 models, using the sequence specific version of the models. Further simulation parameters include a timestep of 15.15 fs, Andersen-like thermostat at 20°C, diffusion coefficient 2.5 simulation units, salt concentration set to 1M NaCl, and saving of the configurations is performed every $10^5$ steps.

To simulate the constant pulling rate experimental condition we used 2 harmonic traps with stiffness constant k = 0.57 pN/nm, acting on the center of mass of one nucleotide on the left and right handle respectively. The initial position of the traps was set to correspond with the position of the respective nucleotides. While the position of the left trap remains fixed, the right trap is moved away in the opposite direction with a rate of 0.14 mm/s. Force is then measured as the distance between the position of the trap and the center of mass of the nucleotide it acts on multiplied by the stiffness constant of the trap. The simulation is performed for 15 ms ensuring the hairpin is completely pulled open. We note however that it’s not easy to establish the correspondence between the simulated time and experimental time due to the highly coarse-grained nature of the model, which also had (100x) higher diffusion coefficient of the nucleotides to speed up the simulations. 7 replicas were run per each condition.

Second type of simulations were the constant force simulations. Here we simulated the DDc1 and DDc2 systems under 2, 5 and 10 pN of constant force, applied to the same nucleotides as in the pulling simulations with moving traps. To first equilibrate the systems, we changed the sequence of the invader strand, keeping the toehold bound to the hairpin and replacing the invader sequence with a poly thymine sequence. The equilibration was performed until the end-to-end distance between the end nucleotides of the respective handle strands to which the force was applied stopped changing (for at least 2 ms). Further on, 8 replicas simulating the invasion process were run per each condition. For all conditions but DDc2 10 pN, 50000 states were recorded corresponding to ~ 76 ms of simulated time. For DDc2 10 pN every replica recorded at least 1000 states corresponding to at least 15 ms simulated time.

##### Free-energy profiles

To obtain the free-energy profiles of DNA systems under different tensions, we simulated a simpler three-strand system based on DNA constructs sequence. We use Virtual Move Monte Carlo (VMMC) algorithm with umbrella sampling. As the full system with DNA handles used in experiments is too large for the VMMC algorithm to sample, we have used a truncated system consisting of three strands (invader, substrate, incumbent, shown in Figure S10). Following the approach from ref[19], we sample the free-energy landscape in terms of the numbers of bonds between the invader strand and substrate and between the incumbent strand and the invader strand. We used umbrella sampling potential with iteratively adjusted weights to sample different state in terms of bonds between the invaders / incumbent strands and the substrate. Weighted Histogram Analysis Method (WHAM) has been used to combine simulations where the sampled system was split into multiple different windows, we have explored systems where the substrate strand has been subjected to a constant force. Opposing forces of magnitude 1, 2, 5, and 10 pN have been applied to a pair of nucleotides in the system (shown schematically in Fig. 3).

### Supplementary Tables and Figures

The values in brackets next to the mean value represent the standard error of the mean (s.e.m.) if not stated otherwise. All errors represent 1σ. If the first digit of the error is one or two, then two significant digits of the error value are given. Otherwise, one significant digit is given.

Table S1: Unfolded nucleotides of DNA and RNA toehold hairpin without trigger strand from constant velocity (CV) pulling and passive mode (PM) experiments (DNA CV: N = 8 cycles, 1 molecule; DNA PM: 3 force steps (in total ≈ 380 s time trace), 1 molecule; RNA CV: N = 8 cycles, 1 molecule; RNA PM: 2 force steps (in total ≈ 150 s time trace), 1 molecule).

| **Nucleic acid** | **Measurement mode** | **Intermediate states (unfolded nucleotides)** | | | | |
|---|---|---|---|---|---|---|
| | | **Fol** | **I1** | **I2** | **I3** | **Unf** |
| DNA | CV | 2 | 18.1 (0.7) | 50 (4) | 60 (4) | 110 |
| | PM | 2 | 17.373 (0.006) | 47.21 (0.13) | 58.60 (0.22) | 110 |
| RNA | CV | 2 | 19.0 (0.9) | 52 (4) | 62 (5) | 110 |
| | PM | 2 | 17.7 (0.6) | 49.0 (1.2) | 59.6 (2.8) | 110 |

For the schematics, the weighted average of CV and PM data were used, divided by two to get unfolded base pairs and rounded to the nearest whole number giving 9, 24 and 29 unfolded base pairs for I1, I2 and I3 for DNA and 9, 25 and 30 for RNA. The number of unfolded nucleotides were calculated from the contour length as described in SI "Stretch and relax cycles". The first base pair was assumed already opened because of fraying[3] (2 unfolded nucleotides for the folded state in Table S1). The total unfolded contour length was in good agreement with an unfolding of 110-2=108 nucleotides assuming a conversion of 0.6 nm/nt for ssRNA and 0.59 for nm/nt for ssDNA, considering fraying and taking the diameter of the nucleic acid into account.

Table S2: Free energies of intermediates compared to NUPACK (DNA, RNA (experiment): N = 1 force step, errors represent statistical errors from population probabilities).

| **Nucleic acid** | **-ΔG$_0$ between intermediate states [$k_BT$] at 25 °C** | | | |
|---|---|---|---|---|
| | Fol <-> I1 | I1 <-> I2 | I2 <-> I3 | I3 <-> Unf |
| DNA (experiment) | 15.236 (0.014) | 30.777 (0.016) | 10.798 (0.025) | 35.96 (0.21) |
| DNA (NUPACK) | 22.2 | 38.9 | 10.6 | 47.8 |
| DNA (mFold) | 22.1 | 37.8 | 11.4 | 47.6 |
| DNA (RNAstructure) | 21.9 | 37.6 | 12.1 | 46.9 |
| RNA (experiment) | 24.075 (0.024) | 50.90 (0.09) | 15.06 (0.23) | N.A. |
| RNA (NUPACK) | 30.9 | 64.5 | 15.4 | 66.3 |
| RNA (RNAstructure) | 31.5 | 59.5 | 17.4 | 70.5 |
| RNA (RNAsoft BLstar) | 26.8 | 51.6 | 15 | 58.9 |
| RNA (Vienna RNA at 37°C) | 26.8 | 51.8 | 15.2 | 58.9 |

The systematic deviations from the predicted values from NUPACK can be explained by systematic errors in the force calibration. We could contribute ~ 10% deviation to an underestimation of the viscosity of the buffer and scavenging system mix. A control hairpin was measured at a custom-built optical tweezers setup[19] with an advanced calibration method[20] and yielded ~ 20 % higher free energies and was within 1% of the predicted NUPACK free energy. This method does not require assumptions on the viscosity and bead radius which allows for a more precise force calibration and explains the deviations observed here. The value for the transition between I3 and Unf is not available since the transition was too slow to observe multiple times in the time span of the experiment.

Table S3: Opened contour length in TMSD processes (DD: N = 6 cycles, 1 molecule; RR: N = 6 cycles, 2 molecules; DDc1: N = 7 cycles, 1 molecule; DDc2: N = 5 cycles, 1 molecule; RRc1: N = 7 cycles, 1 molecule).

| **Hairpin + trigger system** | **Intermediate states (opened contour length [nm])** | | | | | | |
|---|---|---|---|---|---|---|---|
| | TB | IM | IM (theo. 18 bm steps) | FI | FI (theo. 36 bm steps) | FU | FU (theo. 36 bm steps + 38 nts unfolding) |
| DD | 0 | N.A. | N.A. | 32.6 (0.3) | 33.5 | 53.42 (0.24) | 54.9 |
| RR | 0 | N.A. | N.A. | 32.5 (0.3) | 31.7 | 53.5 (0.4) | 53.3 |
| DDc1 | 0 | 16.0 (0.4) | 16.7 | 32.20 (0.26) | 33.5 | 53.09 (0.19) | 54.9 |
| DDc2 | 0 | 16.1 (0.3) | 16.7 | 32.4 (0.6) | 33.5 | 53.1 (0.6) | 54.9 |
| RRc1 | 0 | 14.71 (0.16) | 15.8 | 28.81 (0.08) | 31.7 | 48.83 (0.06) | 53.3 |

Here, the additional opened contour length because of branch migration is listed. The parameters for calculating the theoretical values expected for an opening of this many branch migration steps (bm steps) and/or unfolded nucleotides is written in the SI ("Stretch and relax cycles"). The mean values from the fits are close to the theoretical values within less than 10 %. Only the statistical error is considered here. Systematically slightly lower values of the calculated means can be explained by possible fraying of one base pair at the branch migration junction and drift of the position of the laser beams in the optical tweezer's setup. Also, assumptions on the fixed parameter values $p_{ssRNA}$, $p_{ssDNA}$, $L_{ssRNA/nt}$, $L_{dsRNA/bp}$, $L_{ssDNA/nt}$ and $L_{dsDNA/bp}$ can attribute to the systematic deviations.

Table S4: Opened contour length in TMSD processes with branch migration intermediates (BI) (RRc2: N = 6 cycles, 1 molecule; RRp2 (IM): N = 19 cycles, 1 molecule; RRp2 (FI, FU): N = 7 cycles, 1 molecule).

| **Hairpin + trigger system** | **Intermediate states (opened contour length [nm])** | | | | | | | |
|---|---|---|---|---|---|---|---|---|
| | TB | BI | IM | IM (theo. 18 bm steps) | FI | FI (theo. 36 bm steps) | FU | FU (theo. 36 bm steps +38 nts unfolding) |
| RRc2 | 0 | 11.97 (0.25) | 15.85 (0.17) | 15.8 | 30.3 (0.3) | 31.7 | 50.2 (0.4) | 53.3 |
| | TB | IM | BI | N.A. | FI | FI (theo. 36 bm steps) | FU | FU (theo. 36 bm steps +38 nts unfolding) |
| RRp2 | 0 | 0 | 5.6 (0.4) | N.A. | 29.31 (0.08) | 31.7 | 49.58 (0.18) | 53.3 |

Overall good agreement with the theoretical values and small systematic deviations towards lower values compared to the theoretical ones can be explained as for the errors in Table S3. From the difference in the contour length of the IM to BI state in RRc2 one gets 13.6 branch migration steps starting from the TB state for the position of the BI state. This value was rounded to 14 and is shown in the schematic of Figure S3. The contour length of the branch migration intermediate BI in RRp2 correspond to 9.3 branch migration steps. From a different method in which constant velocity traces of RRp2 were contour length transformed[8] and histograms fitted with a double Gaussian yielded a prominent intermediate with 7.91 (0.09) branch migration steps and a less populated intermediate with 14.62 (0.29) branch migration steps. The value from the constant velocity fits can be understood as an average value of the two intermediates as it was not possible to distinguish between them with this analysis method. The opened contour length of state IM of RRp2 is 0 since the mismatches are located directly after the toehold domain of the trigger strand. Hence, state TB and IM are the same for RRp2.

Table S5: Extrapolated rates and free energies at zero load for DDc1, DDc2 and RRc1. The contour length distances to the transition state $\Delta L_{C,IM\text{-}TS}$ and $\Delta L_{C,TS\text{-}FI}$ are determined by fitting it to a model explained in 'Model for transition rate-extrapolation' (DDc1 (except $-\Delta G_0$): N = 102 force steps, 9 molecules; ($-\Delta G_0$): N = 27 force steps, 9 molecules; DDc2 (except $-\Delta G_0$): N = 60 force steps, 6 molecules; ($-\Delta G_0$): N = 18 force steps, 6 molecules; RRc1 (except $-\Delta G_0$): N = 26 force steps, 3 molecules; ($-\Delta G_0$): N = 9 force steps, 3 molecules).

| Hairpin + trigger system | $\Delta L_{C,IM\text{-}TS}$ [nm] | $\Delta L_{C,TS\text{-}FI}$ [nm] | $k_0^{inv}$ [1/s] | $k_0^{re}$ [1/s] | $-\Delta G_0$ [$k_BT$] | $-\Delta G_0$ (Nupack) [$k_BT$] |
|---|---|---|---|---|---|---|
| DDc1 | 8.4 (0.3) | 8.76 (0.14) | 1.75 (0.16) | 780 (30) | 6.56 (0.11) | 7.94 |
| DDc2 | 8.6 (0.4) | 8.7 (0.4) | 0.052 (0.008) | 1480 (220) | 9.76 (0.10) | 10.01 |
| RRc1 | 7.0 (0.9) | 8.4 (0.9) | 0.009 (0.003) | 230 (90) | 8.64 (0.14) | 11.8 |

For the contour lengths and rates, a weighted average was applied since several molecules with a different amount of constant distance steps and, therefore, different individual errors were used for this analysis. The contour length differences from IM to FI are in good agreement (< 5% deviation) with the theoretically expected values assuming 18 branch migration steps from IM to FI (compare values in Table S4 (IM (theo. 18 bm steps))). The position of the transition state is relatively symmetric between IM and FI with a weak asymmetry for RRc1 (7.0 vs. 8.4). The values are within 18 % of the prediction by the nearest neighbor model NUPACK for the DNA mismatches and 27% for the RNA mismatch for comparable salt and temperature conditions. The systematic deviation of our experimental numbers to lower values as compared to the prediction are likely due to systematic errors of our force calibration (see more details in SI, paragraph after Tab. S2).

Table S6: Opened contour length in DNA-RNA hybrid TMSD (RD1: N = 11 cycles, 3 molecules; RD2: N = 10 cycles, 3 molecules; RD3: N = 6 cycles, 2 molecules; FI: 6 cycles, 2 molecules; FU: N = 3 cycles, 3 molecules).

| Hairpin + trigger system | Intermediate states (opened contour length [nm]) | | | | | | | |
|---|---|---|---|---|---|---|---|---|
| | TB | RD1 | RD2 | RD3 | FI | FI (theo. 36 bm steps) | FU | FU (theo. 36 bm steps + 38 nts unfolding) |
| RD | 0 | 6.71 (0.17) | 14.82 (0.24) | 21.3 (0.3) | 28.6 (0.4) | 32.4 | 49.67 (0.03) | 54 |

The values of TB, RD1, RD2, RD3 and FI were extracted from passive mode experiments because the state RD2 and RD3 were harder to distinguish in the constant velocity experiments. The FU state was then calculated from the average of the FI state and the last unfolding step of the remaining hairpin after full invasion from constant velocity experiments. Systematic deviations towards lower values compared to the theoretical ones of ~ 10 % can be explained as for the deviations in Table S3 and S4. For this reason, the number of branch migration steps completed corresponding to the intermediates RD1, RD2 and RD3 were calculated as the ratio of the contour length from one state and the contour length of the FI state times the total number of branch migration steps e.g. (6.71 nm / 28.6 nm) x 36 bms = 8.45 bms. The values of the three intermediates were rounded to the nearest whole number and are shown in Figure 5.

Table S7: Extrapolated rates and free energies at zero load for RNA-DNA hybrid TMSD. The contour length distances to the transition state are determined by fitting it to a model explained in SI "Model for transition rate-extrapolation (TB <-> RD1 (except $-\Delta G_0$): N = 15 force steps, 3 molecules; ($-\Delta G_0$): N = 15 force steps, 3 molecules; RD1 <-> RD2 (except $-\Delta G_0$): N = 4 force steps, 1 molecule; ($-\Delta G_0$): N = 6 force steps, 3 molecules; RD2 <-> RD3 (except $-\Delta G_0$): N = 4 force steps, 1 molecule; ($-\Delta G_0$): N = 6 force steps, 3 molecules; RD3 <-> FI (except $-\Delta G_0$): N = 4 force steps, 1 molecule; ($-\Delta G_0$): N = 6 force steps, 3 molecules). The errors in the brackets except for the free-energy values represent the standard deviation of the fitting parameter.

| Hairpin + trigger system | Transition(i <-> j) | $\Delta L_{C,i\text{-}TS}$ [nm] | $\Delta L_{C,TS\text{-}j}$ [nm] | $k_0^{ij,inv}$ [1/s] | $k_0^{ji,re}$ [1/s] | $-\Delta G_0$ [$k_BT$] |
|---|---|---|---|---|---|---|
| RD | TB <-> RD1 | 4.2 (1.6) | 3.8 (1.8) | 1.4 (1.2) | 34000 (36000) | 8.34 (0.15) |
| RD | RD1 <-> RD2 | 4.7 (5.7) | 3.7 (9.3) | 0.05 (0.15) | 15000 (82000) | 12.9 (0.4) |

| | | | | | | |
|---|---|---|---|---|---|---|
| RD | RD2 <-> RD3 | 4.0 (5.3) | 3.3 (8.1) | 1.6 (4.7) | 50000 (23000) | 9.2 (0.5) |
| RD | RD3 <-> FI | 3.3 (10) | 3.6 (3.5) | 5 (29) | 33000 (66000) | 9.54 (0.26) |

The large errors (shown in the brackets) reflect the large extrapolation range from 10 pN to 0 pN with data points in the small range from 8 - 12 pN and low statistics (three molecules for TB-RD1 and one molecule for the other transitions). The contour length differences between states *i* and *j* show that similar to the extracted values from contour length transformed passive mode traces the highest contour length difference is between state RD1 and RD2. Apart from that, the values are in a similar range compared to the passive mode values but because of the high error values less significant compared to the passive mode extracted values.

Table S8: Calculated Forward and backward speeds during constant force simulations.

| | Forward (Fwd) and backward (Bwd) speeds during Branch Migration | | | |
|---|---|---|---|---|
| Force | Avg Fwd speeds (ns/step) | Number of counted events | Avg Bwd speeds (ns/step) | Number of counted events |
| 0 pN | 92 | 795 | 121 | 806 |
| 2 pN | 91 | 311 | 343 | 350 |
| 5 pN | 81 | 31 | 1920 | 20 |
| 10 pN | 58 | 57 | 11044 | 4 |

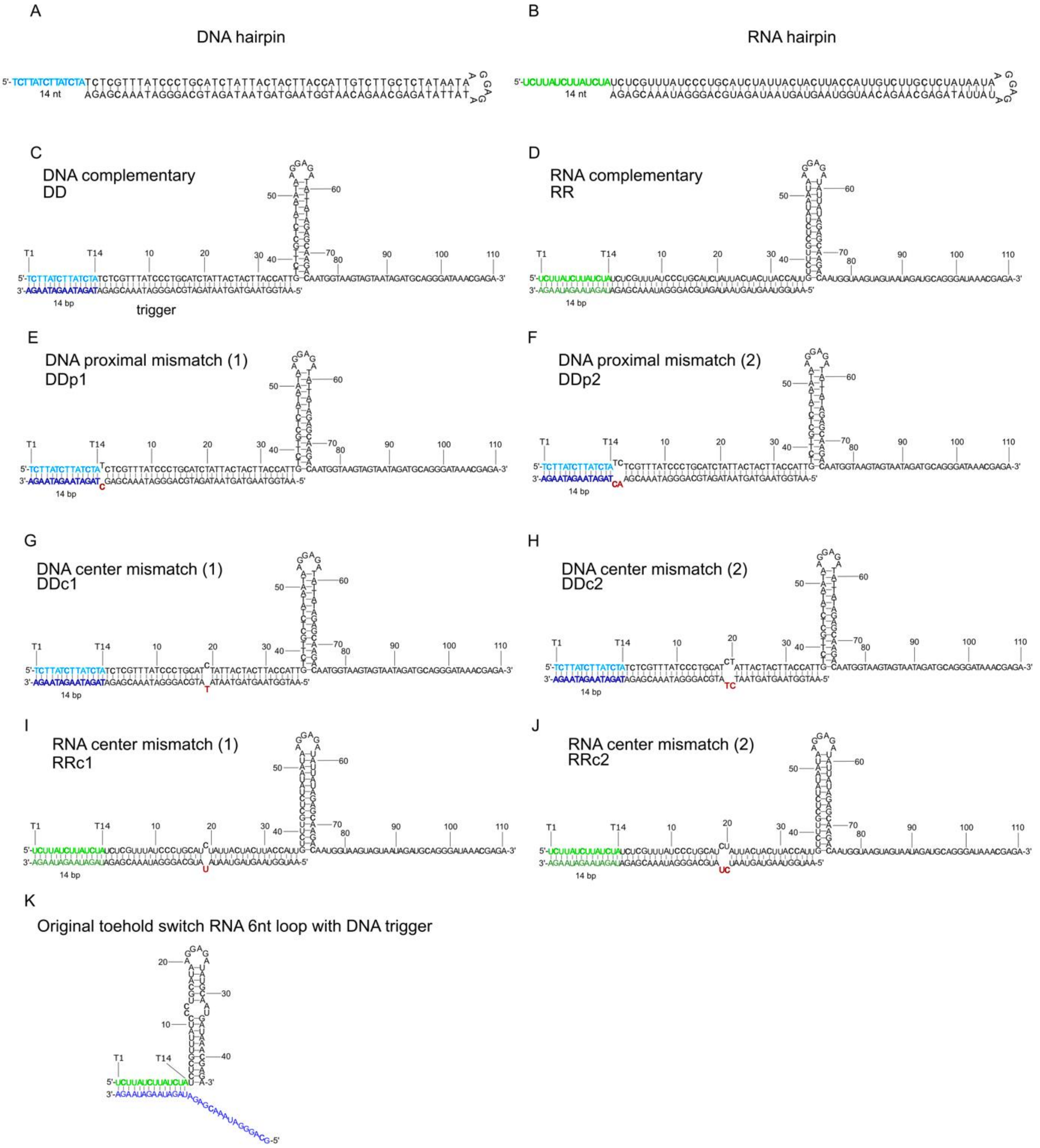

Figure S1. The schematic representation of the toehold hairpins and trigger strands employed in this study, along with their corresponding sequence details. a), b), both the DNA and RNA toehold hairpins are composed of 124 nucleotides, featuring a 14-nucleotide long toehold (T1-T14), a 52-base pair stem, and a 6-nucleotide loop. In specific cases, mismatches were introduced at different positions in the trigger strand for investigation. c), d), the fully complementary trigger strand spans 50 nucleotides and invades the toehold hairpin up to position 36. e), k), 1 proximal mismatch scenarios - DDp1 and RRp1 (C→T, indicated in red) at position 1 following the toehold region. f), l), 2 proximal mismatches scenarios - DDp2 and RRp2 (AG → CA, shown in red) at positions 1 and 2, immediately after the toehold sequence (highlighted in light blue and green). g), i), 1 central mismatch scenarios - DDc1 and DDc2 (C→T, indicated in red) at positions 19. h), j), 2 central mismatches scenarios - DDc2 and RRc2 (GA → TC, shown in red) at positions 19 and 20. K), original toehold switch sequence with 6nt loop with DNA trigger.

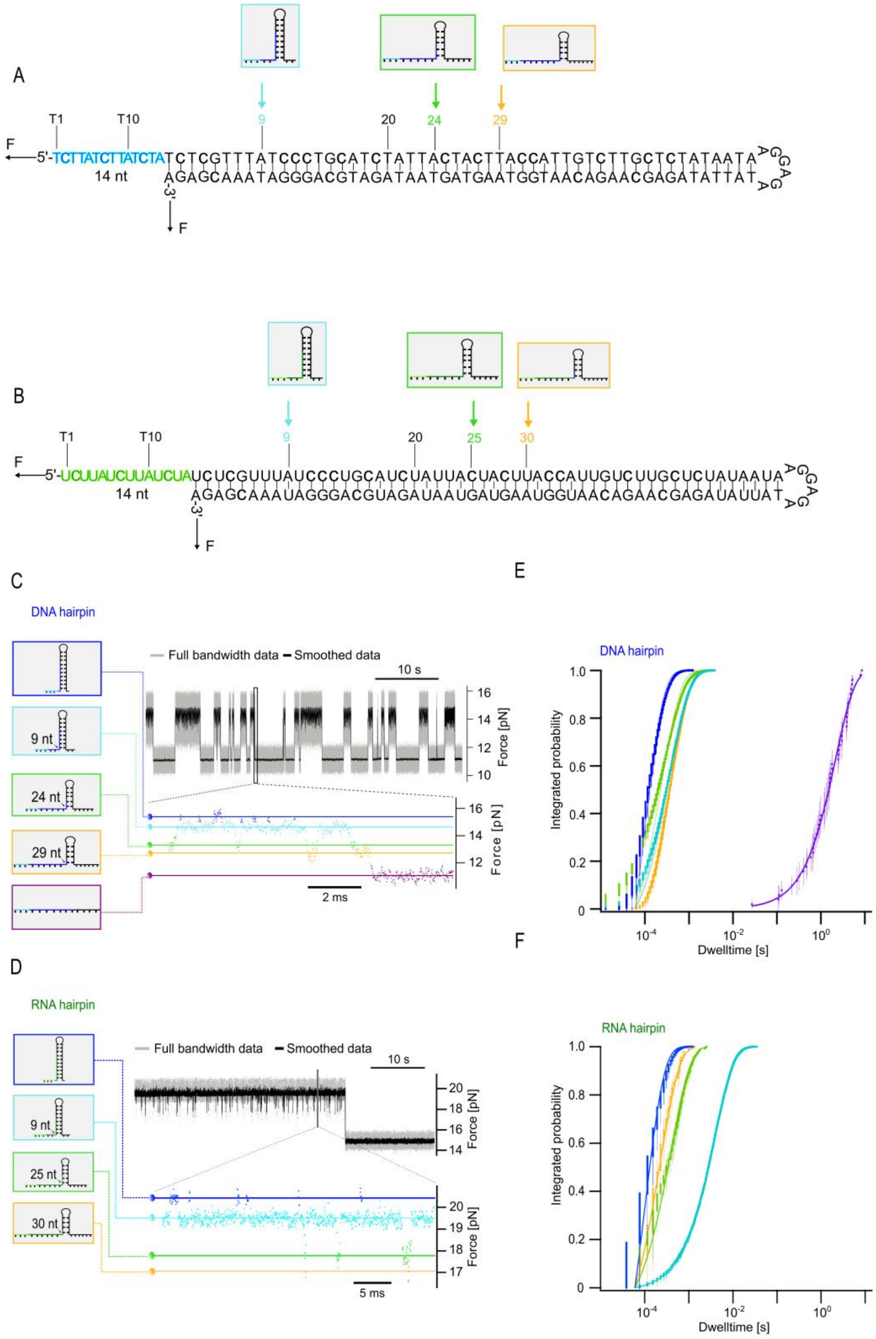

Figure S2. The intermediate states of toehold hairpins from passive mode. A), B), schematic depiction of estimated unzipping intermediate positions for both DNA and RNA toehold hairpins. Each intermediate state is distinguished by different colors. Displayed are force vs. time traces for both DNA and RNA toehold hairpins using passive mode measurements. This approach involves maintaining a constant distance in the laser traps while recording the applied force on the beads. C), D), conformational transitions between intermediate states are evident as shifts between distinct force levels. Each color signifies a specific unzipping intermediate position on the hairpin: folded (blue), intermediate state 1 (light blue), 2 (light green), 3 (orange), and fully opened state (purple). E), F), Dwell time integrated probabilities of each intermediate state derived from the force vs. time traces. In the case of RNA, the dwell times are longer because of the slower kinetics of RNA folding. The slow refolding from the fully unfolded RNA state to the intermediate state shown in orange prevents the observation of multiple fully unfolded dwell times in one constant distance passive mode trace. Therefore, this state is not included in the integrated probability vs. dwell time plot.

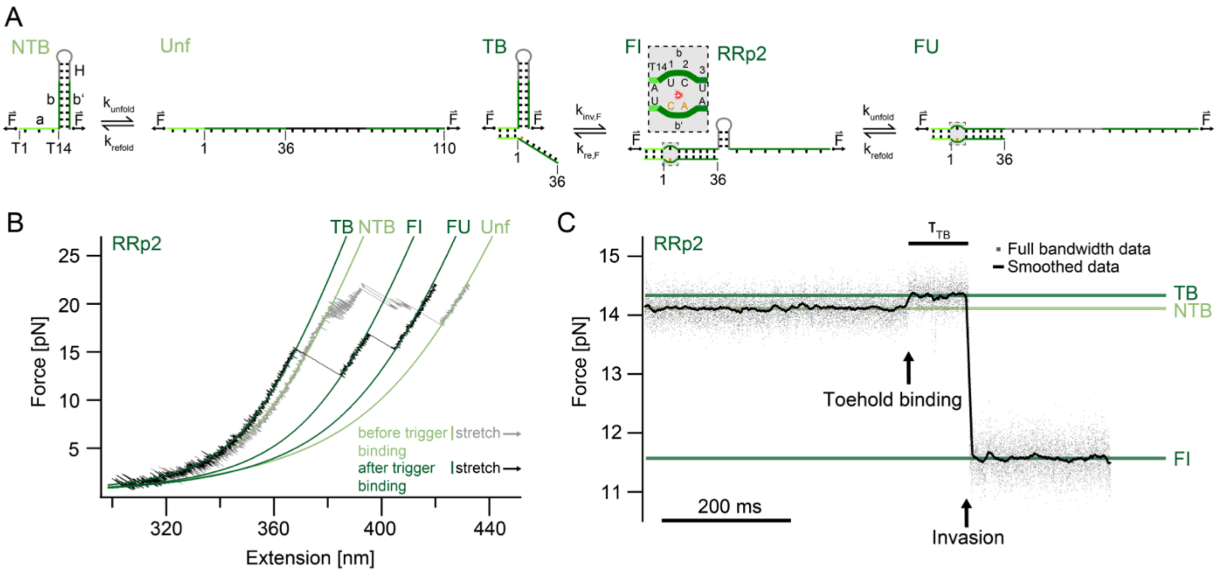

Figure S3. Observation of separate toehold binding and strand invasion. A), Schematic depicting the change in tether extension due to toehold binding, strand invasion and unfolding in the system RRp2. In the absence of a trigger strand in state NTB (non-toehold-bound), the extension of the single-stranded toehold is longer compared to the double-stranded toehold-trigger-bound equivalent in TB (toehold-bound) upon force application. The toehold hairpin transitions from a folded non-toehold bound (NTB) state to an unfolded state (Unf) through unzipping. In the presence of a trigger strand, the toehold-bound state (TB), the fully invaded state (FI) and the fully invaded and unfolded (FU) states are populated in a TMSD process. B), Force-extension traces of the RNA toehold hairpin (gray) before and after binding to an RNA trigger (RRp2) (black). The overlayed traces and fits show the different conformations before and after the trigger is bound associated with different amounts of single-stranded and double-stranded parts contributing to the observed extension. C), Force-versus-time trace of the trigger binding to the RNA toehold hairpin and undergoing strand invasion in the system (RRp2). The toehold binding is characterized by an increase in force since the stretched double-stranded toehold-invader complex has shorter extension than the stretched single-stranded toehold. The invasion is delayed because of the proximal mismatches directly after the toehold binding domain. After ≈ 100 ms, the invasion proceeds which is marked by a drop in force as the invader opens the hairpin.

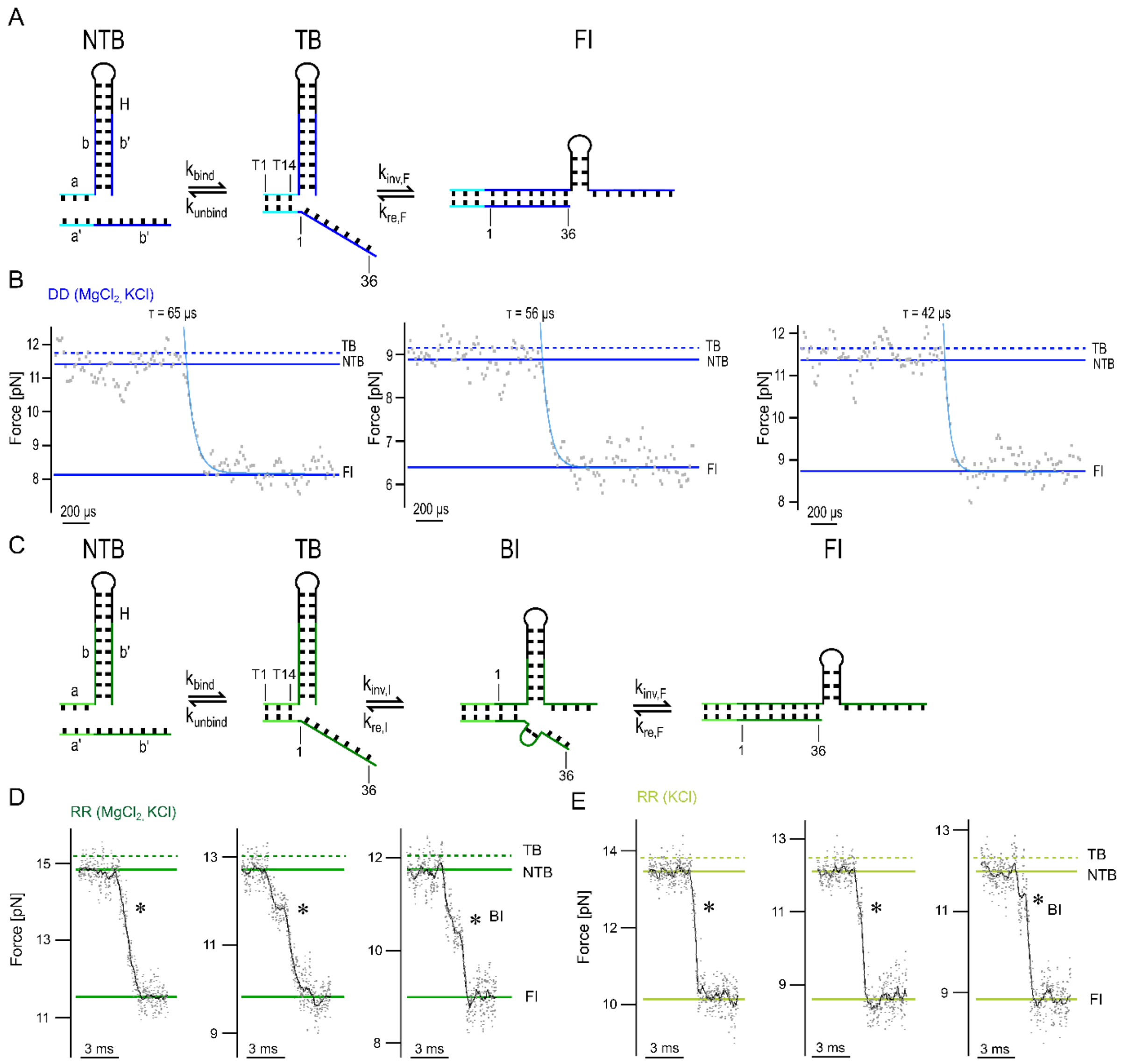

Figure S4. Sample traces of invasion events of toehold hairpins with fully complementary trigger. A), Schematics of the toehold binding and strand invasion of DNA toehold hairpin with fully complementary DNA strand DD, including three different states: non-toehold bound (NTB), toehold bound (TB) and fully invaded (FI). B), Force-versus-time traces of strand invasion of DD while the trap maintains at a constant trap distance under certain salt condition (20 mM $MgCl_2$, 300 mM KCl). Toehold binding and completion of the invasion process occur almost simultaneously and we cannot observe a prolonged lifetime of the toehold-bound state before full invasion as compared to a system like RRp2 (Fig. S3). An estimated TB state at a force higher than the NTB state based on the expected contraction (double-stranded vs. single-stranded, see Discussion “Effect of Mismatches on branch migration kinetics” for details) is marked with dashed lines. The exponential fits show decay times of 65 µs, 56 µs and 42 µs. The data of all events including these three sample traces is summarized in Figure 2D. C), Schematics of the toehold binding and strand invasion of RNA toehold hairpin with fully complementary strand RR, including four different states: non-toehold bound (NTB), toehold bound (TB), branch migration intermediate (BI) and fully invaded (FI). D) E), Force-versus-time traces of strand invasion of RR while the trap maintains at a constant trap distance under certain salt conditions (20 mM $MgCl_2$, 300 mM KCl (D); 300 mM KCl (E)). BI states are shown on the traces with *.

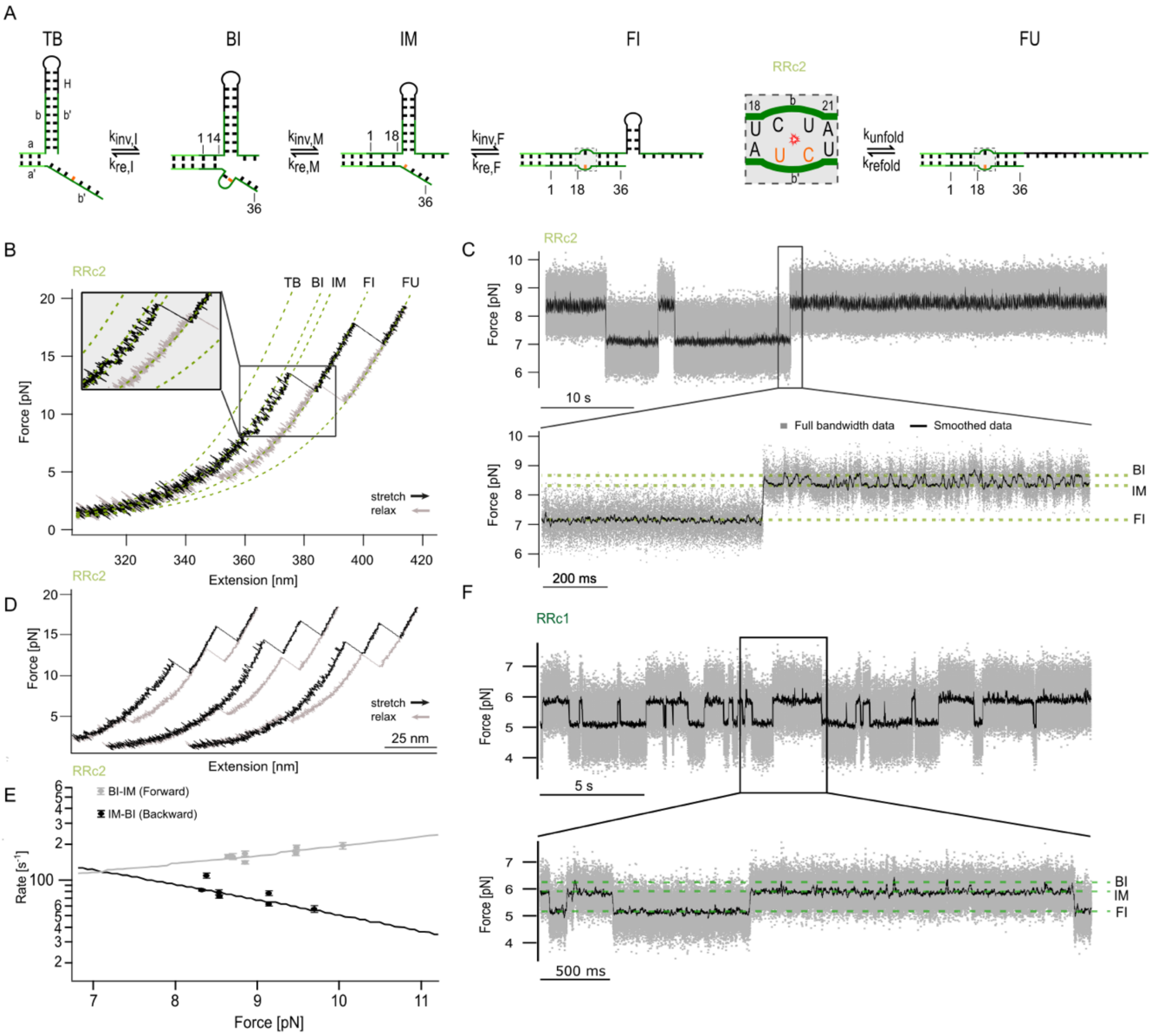

Figure S5. TMSD of RNA toehold hairpin with RNA trigger strand (RRc2). A), Illustration of the RNA toehold hairpin exhibiting 5 distinct states during the force-dependent invading/re-invading and folding/unfolding process triggered by the invading strand: toehold bound (TB), branch migration intermediate before mismatch at position 14 (BI), invasion until mismatch at position 18 (IM), fully invaded (FI), and fully invaded and unfolded (FU). The mismatched base on the trigger strand is highlighted in orange at position 19 and 20. B), Force-extension trace of RRc2. Stretch (black) and relax (gray) cycle using a constant pulling velocity of 0.2 µm/s. Each fit corresponds to a different state as illustrated in A). The transition details between the BI and IM states, influenced by the mismatch and secondary structure in the invader strand, are shown in the inset. C), Force-versus-time traces of the toehold hairpin with the trigger strand (RRc2), maintaining a constant trap distance. Slow transition kinetics are observed between the states IM and FI preventing a quantitative analysis of transition rates as shown in Fig. 3D for systems DDc1, DDc2 and RRc1. D), Additional force-extension traces of the RNA toehold hairpin with the trigger strands (RRc2) showing reproducible transitions between BI and IM. E), extrapolated forward and backward transition rates between the BI and IM states (N = 7 force steps, 1 molecule). F), Force vs. time trace of toehold hairpin with trigger strand RRc1 as comparison showing less prominent excursions from the IM to the BI state compared to RRc2.

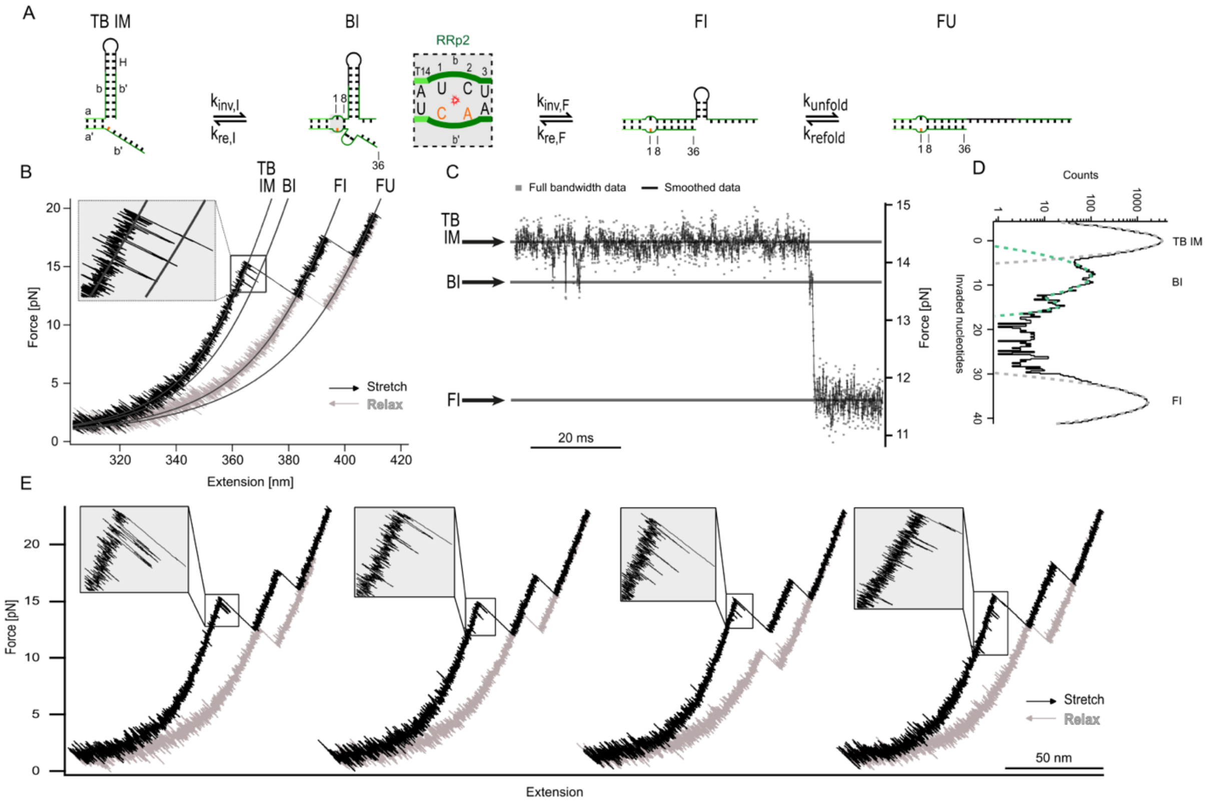

Figure S6. TMSD of RNA toehold hairpin with RNA trigger strand (RRp2). A), Illustration of the RNA toehold hairpin exhibiting 4 distinct states during the force-dependent invading/re-invading and folding/unfolding process triggered by the invading strand: toehold bound and invasion stop before mismatch (TB-IM), branch migration intermediate at position 8 (BI), fully invaded (FI), and fully unfolded (FU). The mismatched base on the trigger strand is highlighted in orange at position 1 and 2. B), Force-extension trace of RRp2. Stretch (black) and relax (gray) cycle using a constant pulling velocity of 0.2 µm/s. Each fit corresponds to a different state as illustrated in A). The transition details between the TB-IM and BI states, caused by secondary structure in the invader strand, are shown in the inset. C), Force-versus-time trace of the invasion event from the TB-IM state to the FI state showing intermediates BI. D), Histogram of invasion data with Gaussian fits of the TB-IM and FI states in gray and of BI in green with one major peak at invaded nucleotide 8 and one minor peak at invaded nucleotide 15 (N = 7 individual histograms from 7 cycles combined into one histogram; 1 molecule). E), Additional force-extension traces of the RNA toehold hairpin with trigger strand showing reproducible transitions between state TB-IM and BI.

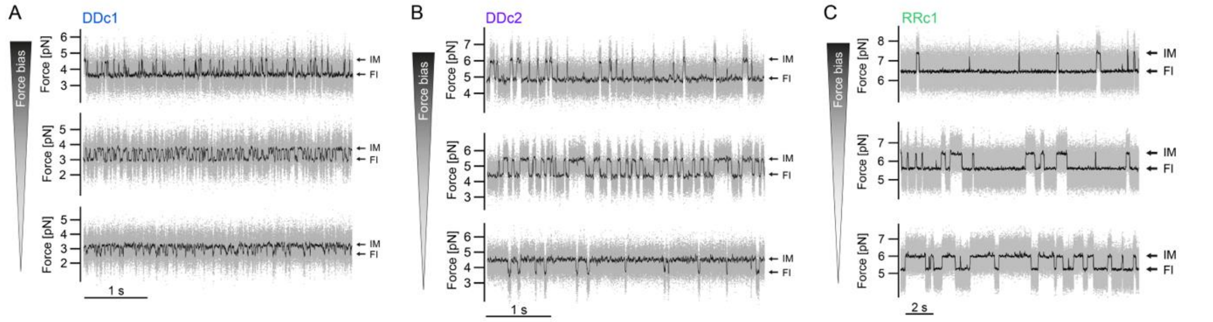

Figure S7. Force versus time traces at different biasing forces for systems DDc1 (A), DDc2 (B) and RRc1 (C). The equilibrium between states IM and FI responds to the applied force, shifting the equilibrium towards the fully invaded state FI with increasing force.

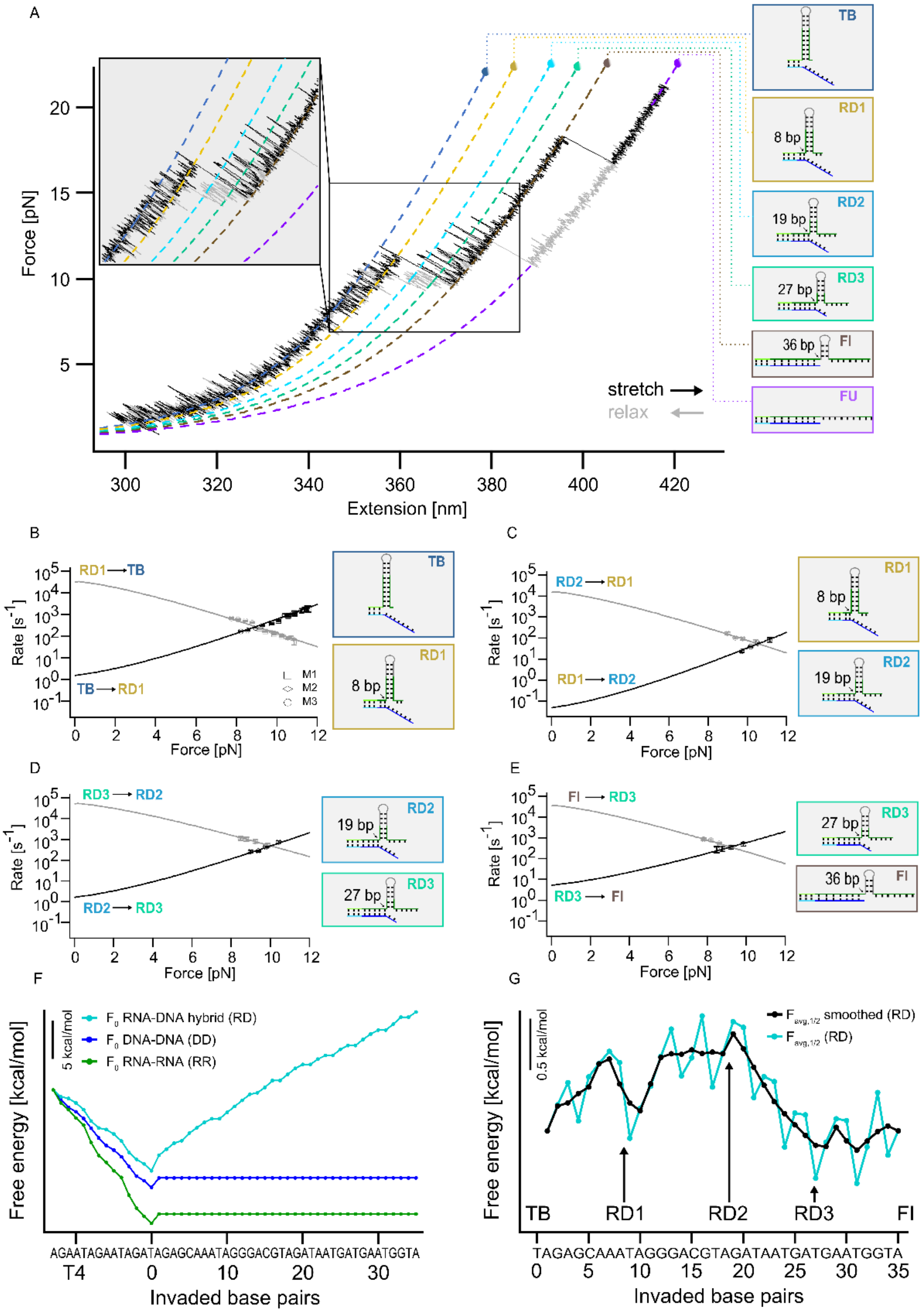

Figure S8. A), Force-extension trace of TMSD of an RNA toehold hairpin with a fully complementary DNA trigger strand (RD). Stretch (black) and relax (gray) cycle using a constant pulling velocity of 0.2 µm/s. Each fit corresponds to a different state as shown in the schematics: Toehold bound (TB), RNA-DNA hybrid intermediates (RD1, RD2, RD3), fully invaded (FI), and fully unfolded (FU). Transition details between state TB and FI are illustrated in the inset. Numbers written in the schematics indicate the opened stem base pairs. B) C) D) E), Force dependence of forward and backward transition rates between each two intermediate states. The black (forward) and gray (backward) lines represent extrapolations of the data based on a theoretical model described in the Supporting Information “Model for transition rate-extrapolation" (B: N = 15 force steps, 3 molecules; C: N = 4 force steps, 1 molecule; D: 4 force steps, 1 molecule; E: 4 force steps, 1 molecule). F), Free-energy profiles for the three strand displacement systems of the branch migration region from our toehold hairpin sequence, calculated using the Nearest Neighbor model. The last invasion (position 36) has no free energy value when the incumbent strand is disattached. The turquoise trace shows an increase in free energy with increasing invaded base pairs as the DNA invader has to overcome the energetic barrier of a more stable RNA base pair. G), Transformed free-energy landscape to $F_{avg,1/2}$ depicting the transition of the DNA invading RNA strand from state TB to FI. The transformation gives the expected landscape at passive mode conditions producing an average force tilting the free energy landscape to an equally likely TB and FI state (see SI "Extracting free energies" for more details on the model used for this transformation). The arrows show the position of the experimentally determined intermediates. RD1 shows a reasonable agreement with a local minimum in the free energy landscape regarding the position.

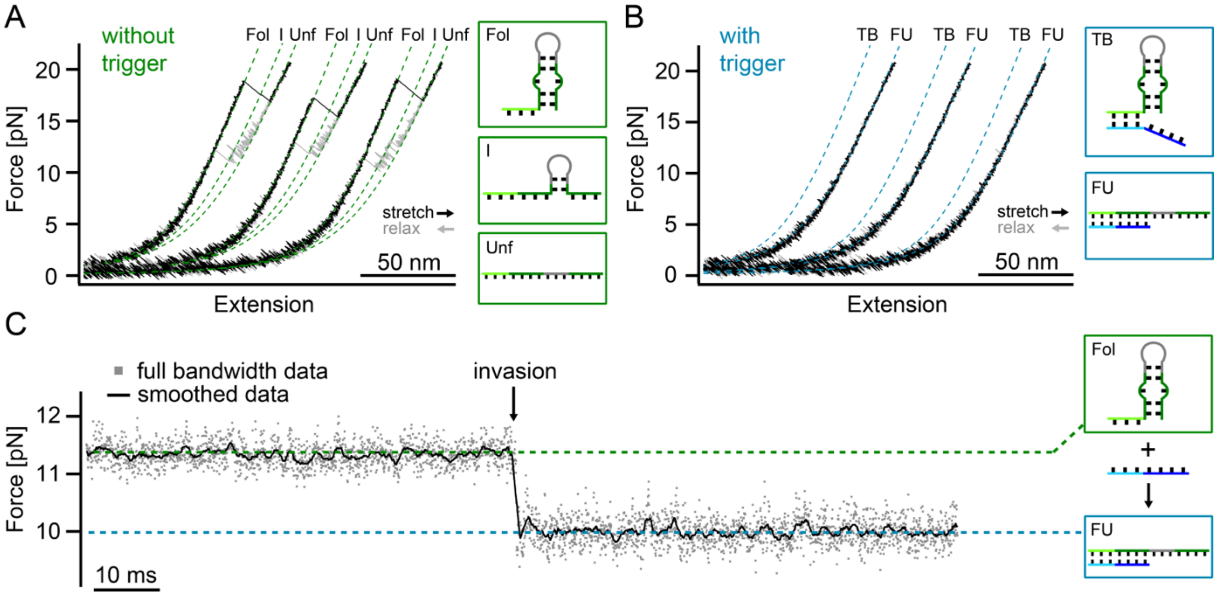

Figure S9. TMSD of an RNA toehold hairpin with a shorter stem and an interior bulge triggered via a DNA trigger using an adapted Toehold Switch design from Green et al[21]. A), B), Force-extension traces before (A) and after (B) trigger binding. Three subsequent stretch (black) and relax (gray) cycles are horizontally shifted with a constant offset for better visualization. Schematics next to the WLC fits represent the different states populated: folded (Fol), intermediate (I), unfolded (Unf), folded and toehold-bound (TB) and fully invaded as well as unfolded (FU). Once the trigger is bound and invaded, no re-invasion events to the TB state are observed, showing that DNA invasion into RNA can shift the equilibrium towards the open state of the toehold hairpin even at very low forces. C), Passive mode trace of the invasion event. Invasion is marked by a drop in force (smoothed trace (black), full bandwidth data (gray)).

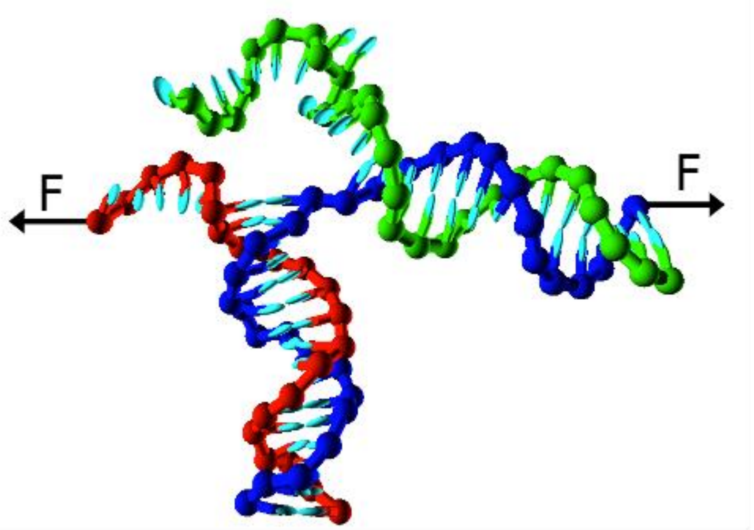

Figure S10. A schematic illustration of a three-strand complex as represented in the oxDNA model. The incumbent (red) is bound to the substrate (blue), and the invading strand is shown in green.  The constant applied force is applied on the nucleotides shown with black arrow.

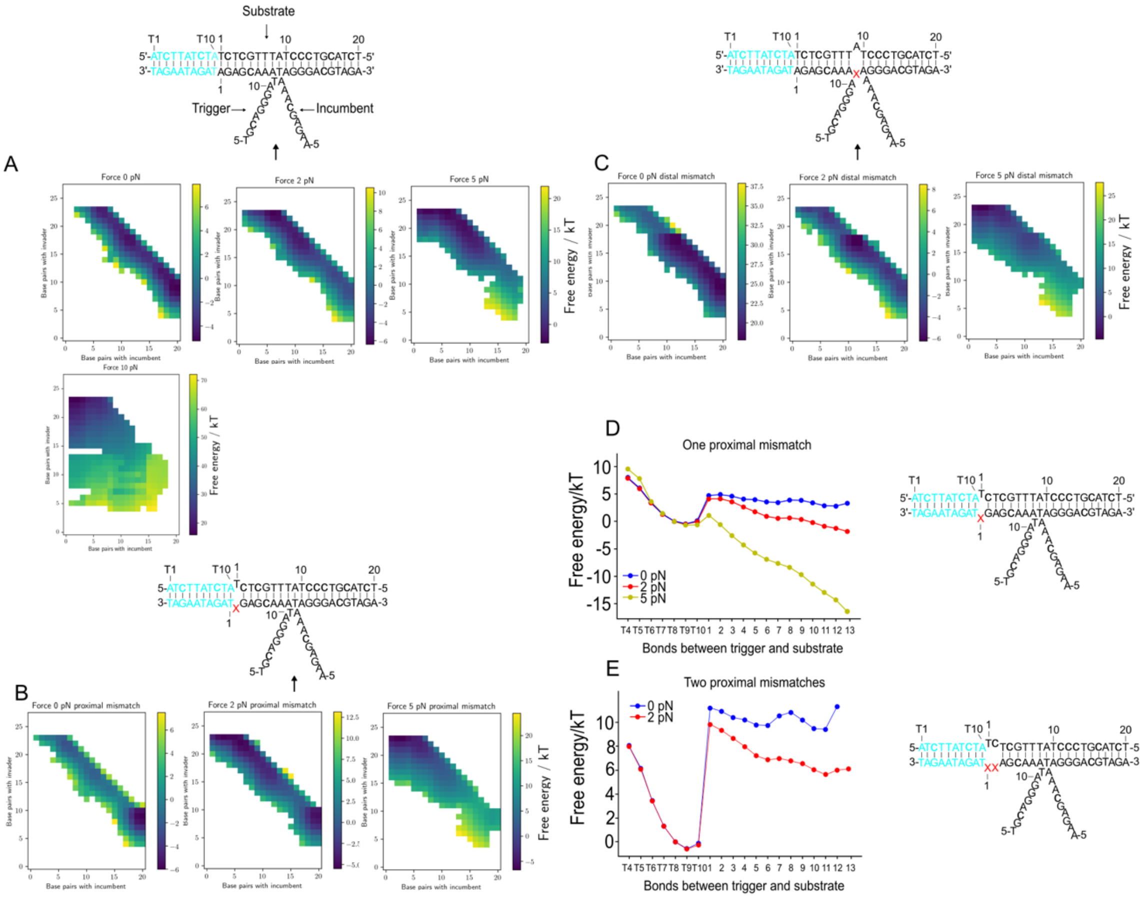

Figure S11. Free energy as a function of bonds between the substrate and the trigger/incumbent strand under different force conditions plotted along the two-dimensional path. A), Free energy landscape of TMSD with a fully complementary trigger under biasing forces 0 pN, 2 pN and 5 pN. B), Trigger has one proximal mismatch. C), Trigger has one distal mismatch. The free energy of the state with the trigger bound to the toehold but with no displacement (trigger base pairs = 10, incumbent base pairs = 20) under each force condition is set to zero. Empty spaces are unsampled. Free-energy profile as a function of number of bases between the incumbent and trigger, shown for D) a trigger that has one mismatch under pulling forces of 0 pN, 2 pN and 5 pN, and E) a trigger that has two proximal mismatches under pulling forces 0 pN and 2 pN.

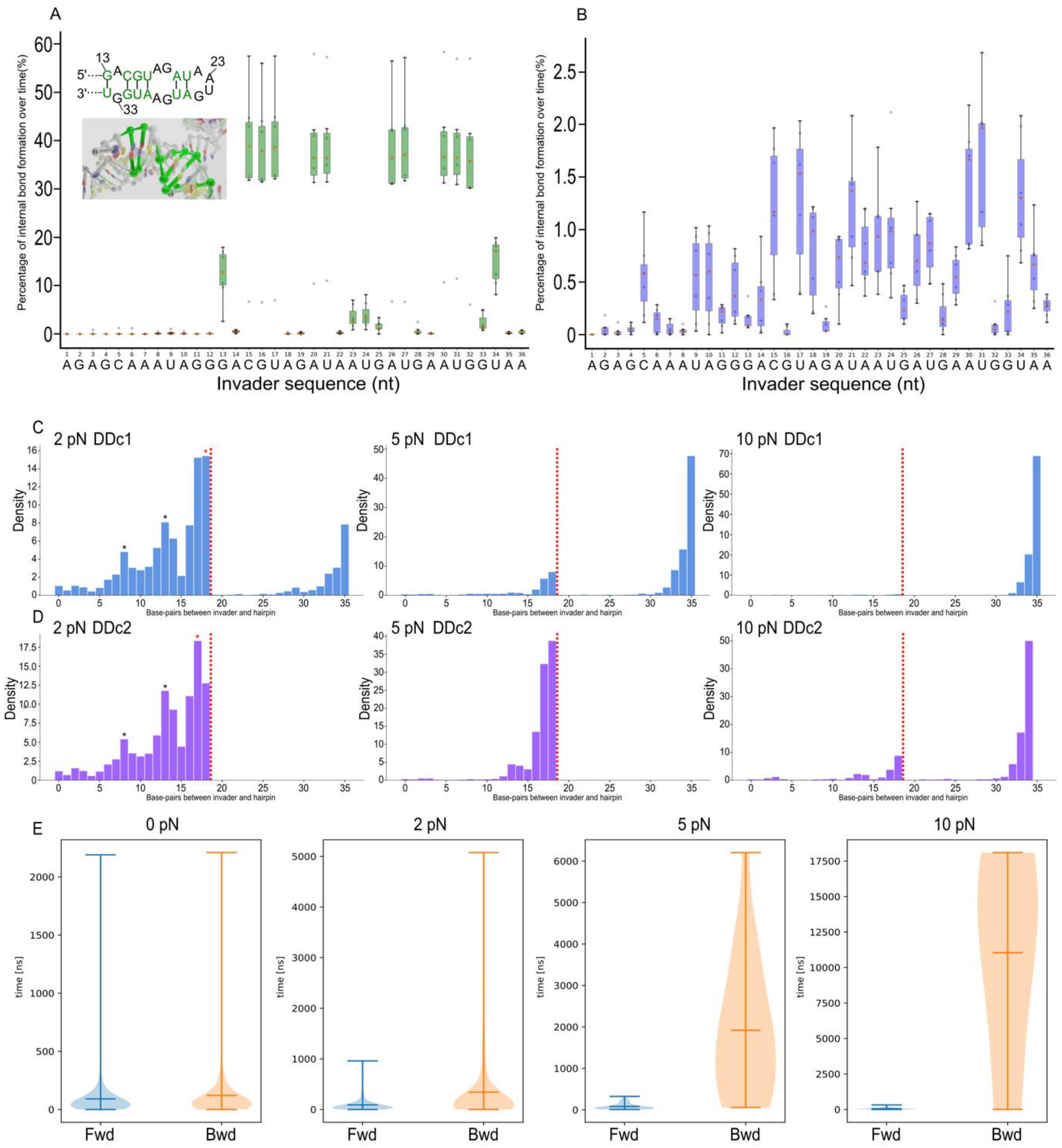

Figure S12. Summary of the oxDNA kinetic simulations. Boxplot of trigger strand bond formation over time for the A) RR and B) DD systems. Obtained from 7 replicas with constant rate pulling at 0.14 mm/s. Inset is showing predicted secondary structure and snapshot of MD trajectory. The potential base pairs are colored in green. C) D) Histograms showing the average percentage of base-pair events for constant force condition simulations (2 pN to 10 pN) of DNA hairpin with invader DDc1 and DDc2 (central mismatch 1 & 2). Each condition has 8 replicas. Stalled intermediate states which arise due to the sequence effect are highlighted with *. Red vertical line corresponds to the position of the mismatch in the trigger strand sequence. E) Violin plot of forward and backward step time obtained from the constant force simulations (DD).

### References

1 Bustamante, C., Marko, J. F., Siggia, E. D. & Smith, S. Entropic elasticity of λ-phage DNA. *Science* **265**, 1599-1600 (1994).

2 Wang, M. D., Yin, H., Landick, R., Gelles, J. & Block, S. M. Stretching DNA with optical tweezers. *Biophysical journal* **72**, 1335-1346 (1997).

3 Woodside, M. T. *et al.* Nanomechanical measurements of the sequence-dependent folding landscapes of single nucleic acid hairpins. *Proceedings of the National Academy of Sciences* **103**, 6190-6195 (2006).
4 Zhang, C. *et al.* The mechanical properties of RNA-DNA hybrid duplex stretched by magnetic tweezers. *Biophysical Journal* **116**, 196-204 (2019).
5 Bloomfield, V. A., Crothers, D. M. & Tinoco, I. Nucleic acids: structures, properties, and functions. *(No Title)* (2000).
6 Lipfert, J. *et al.* Double-stranded RNA under force and torque: similarities to and striking differences from double-stranded DNA. *Proceedings of the National Academy of Sciences* **111**, 15408-15413 (2014).
7 Stigler, J. & Rief, M. Hidden Markov Analysis of Trajectories in Single-Molecule Experiments and the Effects of Missed Events. *ChemPhysChem* **13**, 1079-1086 (2012).
8 Suren, T. *et al.* Single-molecule force spectroscopy reveals folding steps associated with hormone binding and activation of the glucocorticoid receptor. *Proceedings of the National Academy of Sciences* **115**, 11688-11693 (2018).
9 Gebhardt, J. C. M., Bornschlögl, T. & Rief, M. Full distance-resolved folding energy landscape of one single protein molecule. *Proceedings of the National Academy of Sciences* **107**, 2013-2018 (2010).
10 Gebhardt, J. C. M., Clemen, A. E.-M., Jaud, J. & Rief, M. Myosin-V is a mechanical ratchet. *Proceedings of the National Academy of Sciences* **103**, 8680-8685 (2006).
11 Schlierf, M., Berkemeier, F. & Rief, M. Direct observation of active protein folding using lock-in force spectroscopy. *Biophysical journal* **93**, 3989-3998 (2007).
12 Žoldák, G., Stigler, J., Pelz, B., Li, H. & Rief, M. Ultrafast folding kinetics and cooperativity of villin headpiece in single-molecule force spectroscopy. *Proceedings of the National Academy of Sciences* **110**, 18156-18161 (2013).
13 Gardiner, C. (Springer-Verlag, Berlin, 2009).
14 Schink, S. *et al.* Quantitative analysis of the nanopore translocation dynamics of simple structured polynucleotides. *Biophysical journal* **102**, 85-95 (2012).
15 Bohlin, J. *et al.* Design and simulation of DNA, RNA and hybrid protein–nucleic acid nanostructures with oxView. *Nature protocols* **17**, 1762-1788 (2022).
16 Snodin, B. E. *et al.* Introducing improved structural properties and salt dependence into a coarse-grained model of DNA. *The Journal of chemical physics* **142** (2015).
17 Poppleton, E. *et al.* oxDNA: coarse-grained simulations of nucleic acids made simple. *Journal of Open Source Software* **8**, 4693 (2023).
18 Srinivas, N. *et al.* On the biophysics and kinetics of toehold-mediated DNA strand displacement. *Nucleic acids research* **41**, 10641-10658 (2013).
19 Pelz, B. *Enzyme mechanics studied by single molecule force spectroscopy*, Technische Universität München, (2014).
20 Tolić-Nørrelykke, S. F. *et al.* Calibration of optical tweezers with positional detection in the back focal plane. *Review of scientific instruments* **77** (2006).
21 Green, A. A., Silver, P. A., Collins, J. J. & Yin, P. Toehold switches: de-novo-designed regulators of gene expression. *Cell* **159**, 925-939 (2014).
