## Supplementary material for "Single-Molecule Force Spectroscopy of Toehold-Mediated Strand Displacement": SI Sequences and Primers

**Table 1. Sequences of cloning primers and template oligos**

| Primer name | Primer sequence | Description | Annealing temperature (℃) |
|---|---|---|---|
| pt7 DNA linker Fw | gctctagaTAATACGACTCACTATAGGGGCAGGGCTGACGTT | Forward primer binds to DNA linker binding sequence and add pt7 promoter and adds restriction site | 67 |
| Adapter Re | tgcaCTGCAGCACCTGAATTCTGCCCTCAC | Reverse Primer binds to adapter binding sequence and adds restriction site | 66 |
| Trigger complementary Fw | AGATGCAGGGATAAACGAGATAGATAAGATAAGA | Forward primer binds to partial trigger and add the additional sequence based on partial trigger sequence | 58 |
| Trigger proximal mismatches2 Fw | AGATGCAGGGAUAAACGAACUAGAUAAGAUAAGA | Forward primer binds to partial trigger and add the additional sequence with mismatches based on partial trigger sequence | 52 |
| Trigger proximal mismatches1 Fw | AGATGCAGGGAUAAACGAGCUAGAUAAGAUAAGA | Forward primer binds to partial trigger and add the additional sequence with mismatch based on partial trigger sequence | 56 |
| Trigger central mismatches2 Fw | CUAUGCAGGGAUAAACGAGAUAGAUAAGAUAAGA | Forward primer binds to partial trigger and add the | 58 |

| | | | |
|---|---|---|---|
| | | additional sequence with mismatches based on partial trigger sequence | |
| Trigger central mismatches1 Fw | UAUGCAGGGAUAAACGAGAUAGAUAAGAUAAGA | Forward primer binds to partial trigger and add the additional sequence with mismatch based on partial trigger sequence | 58 |
| pt7 Trigger Re | ATTACTACTTACCATTCCTATAGTGAGTCG | Reward primer binds to the pt7 promoter and add partial sequence of the trigger | 58 |
| Trigger Re for expression | GGTTCCATTCTTATCTTATCTA | Primer binds to 3' end of trigger for expression | 50 |

**Table 2. Sequence of toehold DNA hairpins, RNA hairpins and others**

**NOTES: 1. Toehold sequence for RNA expression include pt7 promoter, toehold, DNA linker and adapter binding sequence, dT-Biotin or dT-Digoxigenin**

| DNA sequence | Description | Oligo or Plasmid/ Origin/Resistance |
|---|---|---|
| GGCAGGGCTGACGTTTAACCAGACCAGCGAGTCGTCTTATCTTATCTATCTCGTTTATCCCTGCATCTATTACTACTTACCATTGTCTTGCTCTATAATAAGGAGATATTATAGAGCAAGACAATGGTAAGTAGTAATAGATGCAGGGATAAACGAGAAGTGAGGGCAGAATTCAGGTG | Toehold DNA hairpin with 14nt toehold region and 52 bp stem. DNA linker and adapter binding sequences are located at 5' and 3' end. | oligo |
| GGCAGGGCTGACGTTTAACCAGACCAGCGAGTCGTCTTATCTTATCTATCTCGTTTATCCCTGCATCTATTACTACTTACCATTGTCTTGCTCTATAATAAGGAGATATTATAGAGCAAGACAATGGTAAGTAGTAATAGATGCAGGGATAAACGAGAAGTGAGGGCAGAATTCAGGTG | Toehold DNA hairpin with 14nt toehold region and 52 bp stem and two bulges. DNA linker and adapter binding sequences are located at 5' and 3' end. | oligo |
| TAATACGACTCACTATAGGGGCAGGGCTGACGTTTAACCAGACCAGCGAGTCGTCTTATCTTATCTATCTCGTTTATCCCTGCA | Toehold RNA hairpin expressed under pt7 promoter with 14nt toehold region and | pet28b-modified/ |

| | | |
|---|---|---|
| TCTATTACTACTTACCATTGTCTTGCTC TATAATAAGGAGATATTATAGAGCAAGA CAATGGTAAGTAGTAATAGATGCAGGGA TAAACGAGAAGTGAGGGCAGAATTCAGG TG | 52 bp stem. DNA linker and adapter binding sequences are located at 5' and 3' end. | pBR322 /kanamycin |
| TAATACGACTCACTATAGGGGCAGGGCT GACGTTTAACCAGACCAGCGAGTCGTCT TATCTTATCTATCTCGTTTATCCCTGCA TCTATTACTACTTACCATTGTCTTGCTC TATAATAAGGAGATATTATAGAGCAAGA CAATGGTAAGTAGTAATAGATGCAGGGA TAAACGAGAAGTGAGGGCAGAATTCAGG TG | Toehold RNA hairpin expressed under pt7 promoter with 14nt toehold region and 52 bp stem and two bulges. DNA linker and adapter binding sequences are located at 5' and 3' end. | pet28b-modified/ pBR322 /kanamycin |
| GGCAGGGCTGACGTTTAACCAGACCAGC GAGTCGCACCTGAATTCTGCCCTCACT | Adapter strand which connects the target molecule and DNA handle | oligo |
| GGCGATCTGGTCGTTGATTTGAGTCTGG ATGCGGCCAGATTTGACGAGCAGATGGC CAGAGTCAGGCGTCATTTTTCTGGTACG GAAAGTGATGCGAAAAAAACAGCGGCAG TCGTTGAACAGTCGCTGAGCCGACAGGC GCTGGCTGCACAGAAAGCGGGGATTTCC GTCGGGCAGTATAAAGCCGCCATGCGTA TGCTGCCTGCACAGTTCACCGACGTGGC CACGCAGCTTGCAGGCGGGCAAAGTCCG TGGCTGATCCTGCTGCAACAGGGGGGGC AGGTGAAGGACTCCTTCGGCGGGATGAT CCCCATGTTCAGGGGGCTTGCCGGTGCG ATCACCCTGCCGATGGTGGGGGCCACCT CGCTGGCGGTGGCGACCGGTGCGCTGGC GTATGCCTGGTATCAGGGCAACTCAACC CTGTCCGATTTCAACAAAACGCTGGTCC TTTCCGGCAATCAGGCGGGACTGACGGC AGATCGTATGCTGGTCCTGTCCAGAGCC GGGCAGGGGCAGGGCTGACGTTTAACCA GACCAGCGAGTCG | DNA handle that connects silico beads and target molecule, originally comes from lambda phage sequence with dT-Biotin or dT-Digoxigenin modifications on the 5' end and single strand overhang binding sequence on the 3' end. | PCR product |
| AATGGTAAGTAGTAATAGATGCAGGGAT AAACGAGATAGATAAGATAAGA | DNA trigger strand with fully complementary sequence to the toehold hairpin | oligo |
| AATGGTAAGTAGTAATA***T***ATGCAGGGAT AAACGAGATAGATAAGATAAGA | DNA trigger strand with central one mismatch sequence to the toehold hairpin | oligo |
| AATGGTAAGTAGTAAT***CT***ATGCAGGGAT AAACGAGATAGATAAGATAAGA | DNA trigger strand with central two mismatches sequence to the toehold hairpin | oligo |
| AATGGTAAGTAGTAATAGATGCAGGGAT AAACGAG***C***TAGATAAGATAAGA | DNA trigger strand with proximal one mismatch sequence to the toehold hairpin | oligo |
| AATGGTAAGTAGTAATAGATGCAGGGAT AAACGA***AC***TAGATAAGATAAGA | DNA trigger strand with proximal two mismatches | oligo |

| | | |
|---|---|---|
| | sequence to the toehold hairpin | |
| TAATACGACTCACTATAGGAATGGTAAGTAGTAATAGATGCAGGGATAAACGAGATAGATAAGATAAGA ATGGAACC | RNA trigger strand expressed under pt7 promoter with fully complementary sequence to the toehold hairpin | pet28b-modified/pBR322/kanamycin |
| TAATACGACTCACTATAGG AATGGTAAGTAGTAATA***T***ATGCAGGGATAAACGAGATAGATAAGATAAGA | RNA trigger strand expressed under pt7 promoter with central one mismatch sequence to the toehold hairpin | pet28b-modified/pBR322/kanamycin |
| TAATACGACTCACTATAGG AATGGTAAGTAGTAAT***CT***ATGCAGGGATAAACGAGATAGATAAGATAAGA | RNA trigger strand expressed under pt7 promoter with centrial two mismatches sequence to the toehold hairpin | pet28b-modified/pBR322/kanamycin |
| TAATACGACTCACTATAGG AATGGTAAGTAGTAATAGATGCAGGGATAAACGAG***C***TAGATAAGATAAGA | RNA trigger strand expressed under pt7 promoter with proximal one mismatch sequence to the toehold hairpin | pet28b-modified/pBR322/kanamycin |
| TAATACGACTCACTATAGG AATGGTAAGTAGTAATAGATGCAGGGATAAACGA***AC***TAGATAAGATAAGA | RNA trigger strand expressed under pt7 promoter with proximal two mismatches sequence to the toehold hairpin | pet28b-modified/pBR322/kanamycin |

**Cloning vectors**

| | |
|---|---|
| AGGGATTTTGCCGATTTCGGCCTATTGGTTAAAAAATGAGCTGATTTAACAAAAATTTAACGCGAATTTTAACAAAATATTAACGTTTACAATTTCAGGTGGCACTTTTCGGGGAAATGTGCGCGGAACCCCTATTTGTTTATTTTTCTAAATACATTCAAATATGTATCCGCTCATGAATTAATTCTTAGAAAAACTCATCGAGCATCAAATGAAACTGCAATTTATTCATATCAGGATTATCAATACCATATTTTTGAAAAAGCCGTTTCTGTAATGAAGGAGAAAACTCACCGAGGCAGTTCCATAGGATGGCAAGATCCTGGTATCGGTCTGCGATTCCGACTCGTCCAACATCAATACAACCTATTAATTTCCCCTCGTCAAAAATAAGGTTATCAAGTGAGAAATCACCATGAGTGACGACTGAATCCGGTGAGAATGGCAA | Modified Cloning vector with replicate replication origin pBR322 and Kanamycin resistant. |

| |
|---|
| AAGTTTATGCATTTCTTTCCAGACTTGT<br>TCAACAGGCCAGCCATTACGCTCGTCAT<br>CAAAATCACTCGCATCAACCAAACCGTT<br>ATTCATTCGTGATTGCGCCTGAGCGAGA<br>CGAAATACGCGATCGCTGTTAAAAGGAC<br>AATTACAAACAGGAATCGAATGCAACCG<br>GCGCAGGAACACTGCCAGCGCATCAACA<br>ATATTTTCACCTGAATCAGGATATTCTT<br>CTAATACCTGGAATGCTGTTTTCCCGGG<br>GATCGCAGTGGTGAGTAACCATGCATCA<br>TCAGGAGTACGGATAAAATGCTTGATGG<br>TCGGAAGAGGCATAAATTCCGTCAGCCA<br>GTTTAGTCTGACCATCTCATCTGTAACA<br>TCATTGGCAACGCTACCTTTGCCATGTT<br>TCAGAAACAACTCTGGCGCATCGGGCTT<br>CCCATACAATCGATAGATTGTCGCACCT<br>GATTGCCCGACATTATCGCGAGCCCATT<br>TATACCCATATAAATCAGCATCCATGTT<br>GGAATTTAATCGCGGCCTAGAGCAAGAC<br>GTTTCCCGTTGAATATGGCTCATAACAC<br>CCCTTGTATTACTGTTTATGTAAGCAGA<br>CAGTTTTATTGTTCATGACCAAAATCCC<br>TTAACGTGAGTTTTCGTTCCACTGAGCG<br>TCAGACCCCGTAGAAAAGATCAAAGGAT<br>CTTCTTGAGATCCTTTTTTTCTGCGCGT<br>AATCTGCTGCTTGCAAACAAAAAAACCA<br>CCGCTACCAGCGGTGGTTTGTTTGCCGG<br>ATCAAGAGCTACCAACTCTTTTTCCGAA<br>GGTAACTGGCTTCAGCAGAGCGCAGATA<br>CCAAATACTGTCCTTCTAGTGTAGCCGT<br>AGTTAGGCCACCACTTCAAGAACTCTGT<br>AGCACCGCCTACATACCTCGCTCTGCTA<br>ATCCTGTTACCAGTGGCTGCTGCCAGTG<br>GCGATAAGTCGTGTCTTACCGGGTTGGA<br>CTCAAGACGATAGTTACCGGATAAGGCG<br>CAGCGGTCGGGCTGAACGGGGGGTTCGT<br>GCACACAGCCCAGCTTGGAGCGAACGAC<br>CTACACCGAACTGAGATACCTACAGCGT<br>GAGCTATGAGAAAGCGCCACGCTTCCCG<br>AAGGGAGAAAGGCGGACAGGTATCCGGT<br>AAGCGGCAGGGTCGGAACAGGAGAGCGC<br>ACGAGGGAGCTTCCAGGGGGAAACGCCT<br>GGTATCTTTATAGTCCTGTCGGGTTTCG<br>CCACCTCTGACTTGAGCGTCGATTTTTG<br>TGATGCTCGTCAGGGGGGCGGAGCCTAT<br>GGAAAAACGCCAGCAACGCGGCCTTTTT<br>ACGGTTCCTGGCCTTTTGCTGGCCTTTT<br>GCTCACATGTTCTTTCCTGCGTTATCCC<br>CTGATTCTGTGGATAACCGTATTACCGC<br>CTTTGAGTGAGCTGATACCGCTCGCCGC<br>AGCCGAACGACCGAGCGCAGCGAGTCAG<br>TGAGCGAGGAAGCGGAAGAGCGCCTGAT |

| |
|---|
| GCGGTATTTTCTCCTTACGCATCTGTGC<br>GGTATTTCACACCGCATATATGGTGCAC<br>TCTCAGTACAATCTGCTCTGATGCCGCA<br>TAGTTAAGCCAGTATACACTCCGCTATC<br>GCTACGTGACTGGGTCATGGCTGCGCCC<br>CGACACCCGCCAACACCCGCTGACGCGC<br>CCTGACGGGCTTGTCTGCTCCCGGCATC<br>CGCTTACAGACAAGCTGTGACCGTCTCC<br>GGGAGCTGCATGTGT |
